# Broccoli: combining phylogenetic and network analyses for orthology assignment

**DOI:** 10.1101/2019.12.13.875831

**Authors:** Romain Derelle, Hervé Philippe, John K. Colbourne

## Abstract

Orthology assignment is a key step of comparative genomic studies, for which many bioinformatic tools have been developed. However, all gene clustering pipelines are based on the analysis of protein distances, which are subject to many artefacts. In this paper we introduce Broccoli, a user-friendly pipeline designed to infer, with high precision, orthologous groups and pairs of proteins using a phylogeny-based approach. Briefly, Broccoli performs ultra-fast phylogenetic analyses on most proteins and builds a network of orthologous relationships. Orthologous groups are then identified from the network using a parameter-free machine learning algorithm. Broccoli is also able to detect chimeric proteins resulting from gene-fusion events and to assign these proteins to the corresponding orthologous groups. Tested on two benchmark datasets, Broccoli outperforms current orthology pipelines. In addition, Broccoli is scalable, with runtimes similar to those of recent distance-based pipelines. Given its high level of performance and efficiency, this new pipeline represents a suitable choice for comparative genomic studies.

Broccoli is freely available at https://github.com/rderelle/Broccoli.

## Introduction

Orthologous genes are genes originating from a speciation event, as opposed to paralogous genes originating from a gene duplication event (Koonin 2005). The identification of either orthologous pairs or orthologous groups of genes (i.e. independent sets of orthologs found at a given taxonomic level) is the primary step of most comparative genomic studies, since it provides genetic equivalences between species. For instance, the extrapolation of functional genetic discoveries made from experimental model species to distantly related species, including to humans in medicine and in environmental toxicology, requires a precise mapping of orthologs across species.

Assigning gene orthology across distantly related species typically consists of identifying ancient speciation and gene duplication events from the comparisons of present gene or protein sequences. This task is highly challenging for many reasons. The combination of successive speciation and gene duplication events, with the latter often being associated with gene losses and gene conversions (Kondrashov 2012; Pich and Kondrashov 2014; Harpak, et al. 2017), tends to blur the distinction between orthologs and paralogs. In addition, incomplete lineage sorting (Maddison 1997), and the transfers of genetic material between species (i.e. lateral gene transfers) (Soucy, et al. 2015) and between genes (i.e. gene-fusions) (Zmasek and Godzik 2012), all create complex reticulate gene histories. Finally, the heterogenous evolutionary rate of proteins, with known variations across species and over time (Dorus, et al. 2004; Kawahara and Imanishi 2007), and gene prediction errors (e.g. missing, truncated or fused genes) are also important sources of background noise in orthology inferences.

Current *de novo* clustering algorithms are all based on the analysis of pairwise protein distances. Two main approaches have been proposed: distances can be analysed (i) using the best bi-directional hits (BBH) approach or one of its derivative to infer orthologous pairs as implemented in Hieranoid or OMA (Huynen and Bork 1998; Roth, et al. 2008; Schreiber and Sonnhammer 2013; Sonnhammer and Ostlund 2015; Cosentino and Iwasaki 2019), or (ii) using the Markov Cluster algorithm (MCL) to infer orthologous groups from the network of similarities (Dongen 2000; Li, et al. 2003; Emms and Kelly 2015), orthologous groups that can further be analysed using phylogenetic analyses and a species tree reconciliation approach to infer orthologous pairs (Emms and Kelly 2019). The BBH approach is highly precise but is inclined to miss orthologous pairs due to its highly constrained nature (Dalquen and Dessimoz 2013). By contrast, the MCL approach is generally inclusive but unavoidably merges orthologous groups with high sequence similarity, thus lacks precision. Finally, it is important to note that similarity distances are always an underestimate of the true evolutionary distances due to the saturation of sequences, making it difficult for these distance-based approaches to resolve ancient gene histories.

As an alternative, the use of phylogenetic analyses as a first step has been proposed (Gabaldon 2008). The basic principle of this approach is to build a phylogenetic tree for each protein and its similarity hits, and to infer orthologous relationships based on the taxonomic distribution of hits in the trees. The delineation between orthologs and paralogs is made here from the analysis of phylogenetic relationships rather than protein distances (Huerta-Cepas, et al. 2007; Vilella, et al. 2009; Huerta-Cepas, et al. 2014). The promise of this ‘phylogeny-based’ approach at improving orthology inferences has three important caveats: (i) the many hundreds of thousands of phylogenetic analyses required by this approach must be computationally efficient, (ii) new methods for the delineation of orthologous groups must be proposed and (iii) a phylogeny-based pipeline must be made freely available to the research community.

Here we introduce Broccoli, an open-source pipeline for *de novo* orthology assignment using a phylogeny-based approach. Briefly, Broccoli performs ultra-fast phylogenetic analyses and extracts successively two sets of orthologous relationships from the trees. The first set is used to build an orthology network (as opposed to networks of similarity distances), from which orthologous groups are identified using a label propagation algorithm. Then a more precise second set is defined to identify pairs of orthologous genes within each orthologous group.

The performance of Broccoli was assessed by using a custom benchmark dataset for orthologous groups, and the Quest of Orthologs 2018 benchmark dataset for orthologous pairs (Altenhoff, et al. 2016; Glover, et al. 2019). In these tests, we compared Broccoli to recent distance-based pipelines combined with fast similarity search algorithms (e.g. DIAMOND, MMseq2 (Buchfink, et al. 2015; Steinegger and Soding 2017)) since blastp, which is two orders of magnitude slower, would not be usable for large datasets.

## Materials and methods

Broccoli is a pipeline written in Python 3 that requires the ete3 library (Huerta-Cepas, et al. 2010). It is composed of four steps as summarized in Figure 1A and described below. The rationale of Broccoli is that, since single gene trees are expected to be too inaccurate to directly infer orthology relationships, as many trees as the number of sequences will be inferred (Steps 1 and 2) and orthology will be inferred from the consensus of information extracted from these multiple trees using a network analysis (Step 3).

**Figure 1:**
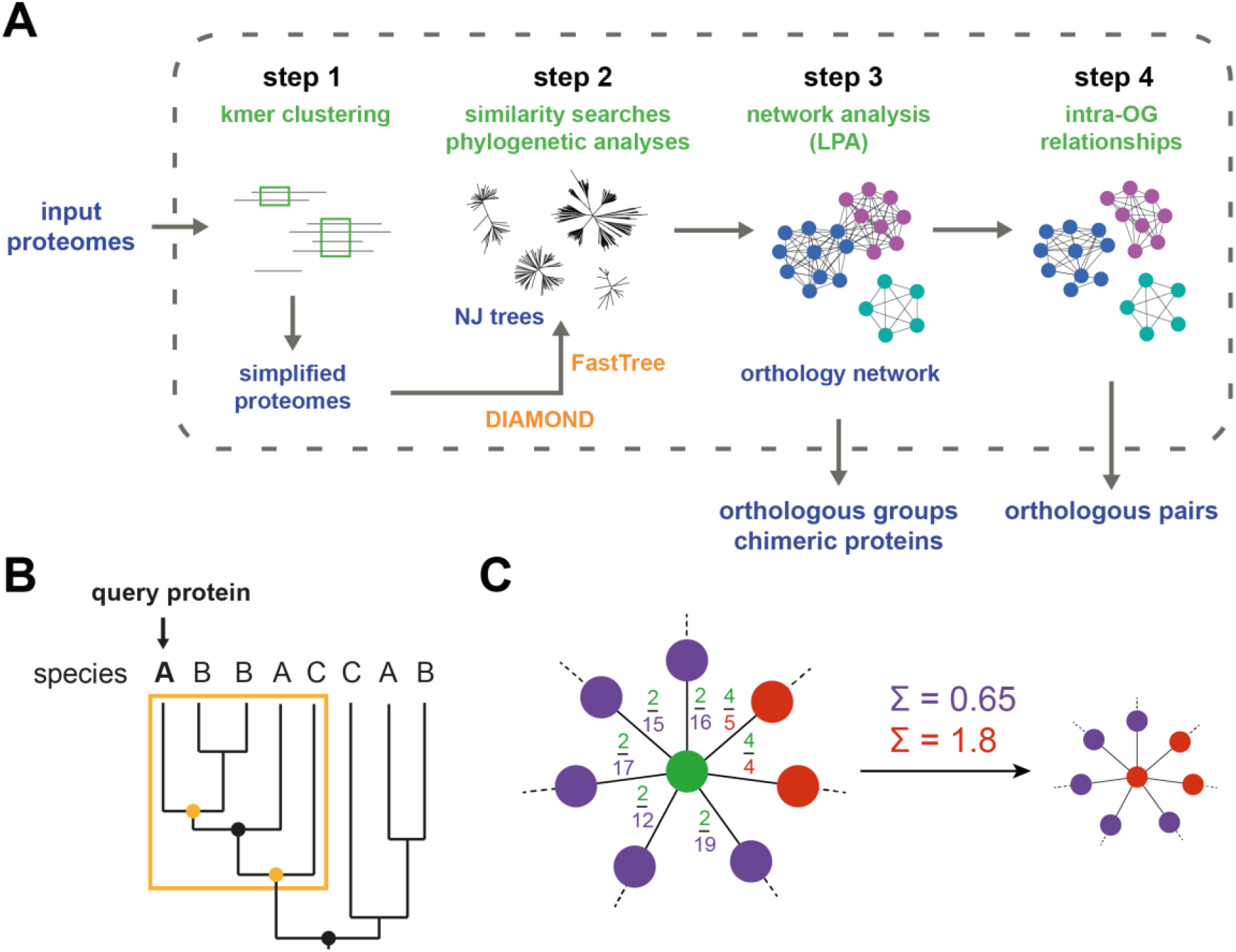
Key aspects of Broccoli. **A**: Overview of the pipeline. Data, external programs and processes are coloured in blue, orange and green respectively. **B**: Example of the species overlap approach on a gene-tree obtained from 3 species A, B and C. The nodes fulfilling the species overlap criteria are indicated by orange dots, and the resulting orthologous group is delineated by the orange rectangle. **C**: Example of the label propagation, with labels represented by colours (green, purple and red). The node with the green label is about to exchange its label with one of its neighbours. The fractions present on each edge represent the weights AB where A is the green node and B its neighbour. The green node takes the red label since the sum of the ‘red weights’ is higher than the sum of the ‘purple weights’.

### Step 1: kmer pre-clustering

The objective is to simplify proteomes without loss of information and therefore to decrease the computational time of steps 2 and 3. Broccoli first converts the protein names into unique identifiers. The proteome of each species is then independently clustered using kmers of amino-acids. For each cluster of sequences, the longest one is retained for further analysis while others are set-aside and will be re-injected into the orthologous groups and orthologous pairs at the corresponding steps. This step aims at reducing the number of proteins to be analysed by removing allelic variants and ‘recent’ duplicates. By default, the kmer size is set to 100 amino-acids. This high value prevents the grouping of paralogs between closely related species. But the kmer size can be reduced when distantly related species are analysed (e.g. species belonging to different eukaryotic supergroups).

### Step 2: similarity searches and phylogenetic analyses

Broccoli then builds a phylome (i.e. the set of gene trees (Huerta-Cepas, et al. 2007)) for each species by comparing its proteins against other proteomes and performing phylogenetic analyses in possible cases of gene duplications (i.e. cases of multiple hits for at least one species). For each simplified proteome, similarity searches against all proteomes are individually performed using DIAMOND under the ‘most-sensitive’ option and the *N* best hits per species are reported (*N* is set to 6 by default). Then, for each query protein, all its hits are considered orthologs to each other if no species have multiple hits (referred thereafter as ‘set-aside orthologous pairs’). Otherwise, the DIAMOND pairwise alignments between the query and each of its target sequences are combined together to build a trimmed alignment by allowing a fraction *g* of missing data per position (*g* is set to 0.7 by default). The trimmed alignment is then analysed using FastTree2 (Price, et al. 2010) to produce a BioNJ tree that is rooted using the midpoint method.

To our knowledge, it is the first time DIAMOND (or blastp) alignments are used to perform phylogenetic analyses instead of classical multiple sequence alignments (MSA). The main advantage of this approach is a considerable decrease of the computational time since alignments are already computed during the similarity searches. But the use of these pairwise alignments also have two additional advantages: (i) only sequence fragments matching the query sequence are used for phylogenetic analyses while MSA, which operate on full length sequences, usually include unaligned blocks that create phylogenetic noise, and (ii) short sequences are often misaligned in MSA but not in pairwise alignments.

### Step 3: identification of orthologous groups

In this third step, Broccoli builds an orthology network from which orthologous groups are isolated using a machine learning algorithm. Broccoli first delineates orthologous groups in each rooted tree using a relaxed ‘species overlap’ approach as defined in (Huerta-Cepas, et al. 2007). Briefly, the trees are traversed from the query protein to the root and, at each node, the taxonomic composition of the two sets of leaves emerging from that node are compared (an example is provided in Figure 1B). The two sets of leaves are considered part of the same orthologous group if (i) there is no common species between the two sets (i.e. no ‘overlap’ species) or (ii) there is only one common species and at least two unique species in both sets (i.e. species not present in the other set). Broccoli identifies the deepest node of the tree fulfilling this ‘species overlap’ criteria, and builds orthologous pairs between all leaves belonging to that node and also paralogous pairs between these leaves and all remaining leaves of the tree.

The orthologous and paralogous pairs extracted from all trees are then combined with the ‘set-aside orthologous pairs’ from Step 2, to build an undirected network of orthologous relationships. An edge between two proteins A and B is formed if they have been identified (i) orthologs at least twice (since at the very least A has been compared to B and B compared to A) and (ii) more often as orthologs than as paralogs. The edge weight w(AB) from A to B is defined as:

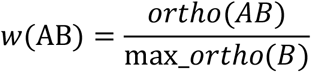

where ‘ortho(AB)’ corresponds to the number of times A and B have been identified as orthologs, and ‘max_ortho(B)’ corresponds to the maximum number of times B has been found to be an ortholog with any other protein. Therefore, the weight w(AB), ranging from infinitesimal to 1, represents the relevance of the orthologous relationships between A and B with respect to the reference node B. The weights are asymmetric since w(AB) might be different from w(BA).

Given the fast phylogenetic analyses, tree rooting and orthologous group delineations performed by Broccoli, the orthology network is expected to be noisy. But one can expect that truly orthologous proteins will be much more often connected and with higher weights among themselves than with paralogous proteins. A label propagation algorithm (LPA; (Raghavan, et al. 2007)) is applied to the orthology network to identify node communities (i.e. orthologous groups). The LPA used here, described in Supplemental Material 1, is asynchronous and weighted (using the asymmetric edge weights described above), resulting in a highly precise community delineation. An example is given in Figure 1C, in which the ‘green’ node is assigned by Broccoli to the ‘red’ community due to the high relevance of its orthologous relationships with this community (an unweighted LPA would assign this node to the ‘purple’ community with which it has more connexions). This algorithm is also fast, with convergence of the labels being reached after only a few generations (Supplemental Material 1) and, in the absence of any random choice, fully deterministic.

Finally, two types of error corrections are applied to the detected communities (i.e. orthologous groups). First, Broccoli attempts to remove spurious hits, which are defined as proteins having less than *n* proteins of the orthologous group in their own similarity hits (*n* is set to 2 by default; connected components of three or less proteins are not subject to LPA and corrections). Proteins considered as spurious hits are then removed from their orthologous group, and therefore from the classification. Second, since proteins are initially assigned to a unique orthologous group, Broccoli aims at detecting gene-fusions and corrects the classification accordingly. Proteins resulting from gene-fusion events are detected among nodes connected to several communities using the method described in Supplemental Material 1. Proteins that are identified as chimeric proteins are then added to all orthologous groups involved in their corresponding fusion event.

### Step 4: identification of orthologous pairs

While orthologous relationships were extracted at Step 3 to delineate orthologous groups, Broccoli builds a new set of orthologous relationships that considers gene duplication events within each orthologous group. The method here is the same as described in Step 3 but with two differences: (i) proteins not belonging to the orthologous group are first removed from the ‘set-aside orthologous pairs’ and from the rooted trees, and (ii) orthologous and paralogous pairs are built at each tested node from the rooted trees – not only at the deepest node fulfilling the species overlap criteria. Finally, for each pair of proteins A and B belonging to this orthologous group, a ratio R(AB) is calculated as

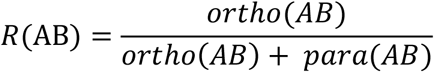

where ‘ortho(AB)’ and ‘para(AB)’ represents the number of times A and B have been found as orthologs and as paralogs respectively. The two proteins will thus be reported as orthologs if their ratio R is superior to a threshold *r* (*r* is set to 0.5 by default).

### Performance tests

The paraBench dataset was built from an in-house collection of phylogenetic markers. The dataset and the performance metrics are fully described in Supplemental Material 2. As opposed to the benchmark of orthologous pairs, the performance metrics were calculated considering all possible pairs within each orthologous group. The dataset, reference clustering, python script to compute the performance metrics and the results obtained in this study are all available at https://github.com/rderelle/paraBench.

In addition, the Quest for Orthologs (QfO) benchmark 2018 dataset was downloaded from the EBI ftp server, then analysed by Broccoli using the ‘not_same_sp’ option and by varying the *r* threshold value using the ‘-ratio_ortho’ option. The resulting sets of orthologous pairs were submitted to the OpenEBench website to run the QfO benchmark.

The versions of the pipelines and programs, and command lines, used in this study are indicated in Supplementary Material 3. All benchmark outputs and QfO raw results are available to download in the Zenodo research data archive at https://zenodo.org/record/3710751.

### Running time analyses

The running time analyses based on fungal datasets were performed using 4 CPUs of an Intel Xeon Gold 6248 processor and 40 GB of RAM memory. The fungal datasets correspond to those used in (Emms and Kelly 2019), with one modification performed on the 64 species dataset (see readme file in the Zenodo archive). All datasets are available to download in the Zenodo research data archive at https://zenodo.org/record/3710751.

The QfO 2018 runtimes were measured using 8 CPUs of the same processor and 60GB of RAM memory.

## Results

### Benchmark of orthologous groups

The quality assessment of orthologous group predictions was performed using a custom-built benchmark dataset (named ‘paraBench’) comprising 17 eukaryotic species and 52 orthologous groups (see Supplemental Material 2). In this benchmark, we compared Broccoli to two recent distance-based pipelines: OrthoFinder2 (Emms and Kelly 2019), which uses the MCL algorithm after distance corrections to mitigate the impact of evolutionary rate differences between species, and Sonicparanoid (Cosentino and Iwasaki 2019), which employs a BBH approach. Broccoli produced the highest recall score value, closely followed by OrthoFinder2, thanks to its distance corrections (Figure 2; Supplemental Material 3). Finally, Sonicparanoid, which over-split orthologous groups due to the stringency of the BBH approach, scored the lowest. On the precision side, Broccoli also performed better than the two other pipelines with a score of 0.973. OrthoFinder2 scored the lowest precision value indicating a tendency to over-merge closely related orthologous groups. Running Sonicparanoid using the ‘most-sensitive’ option as recommended for distantly related species yielded a slightly different protein clustering, yet achieving the same performance metrics. Overall, Broccoli scored the highest on this performance benchmark (F1-score in Figure 2).

**Figure 2:**
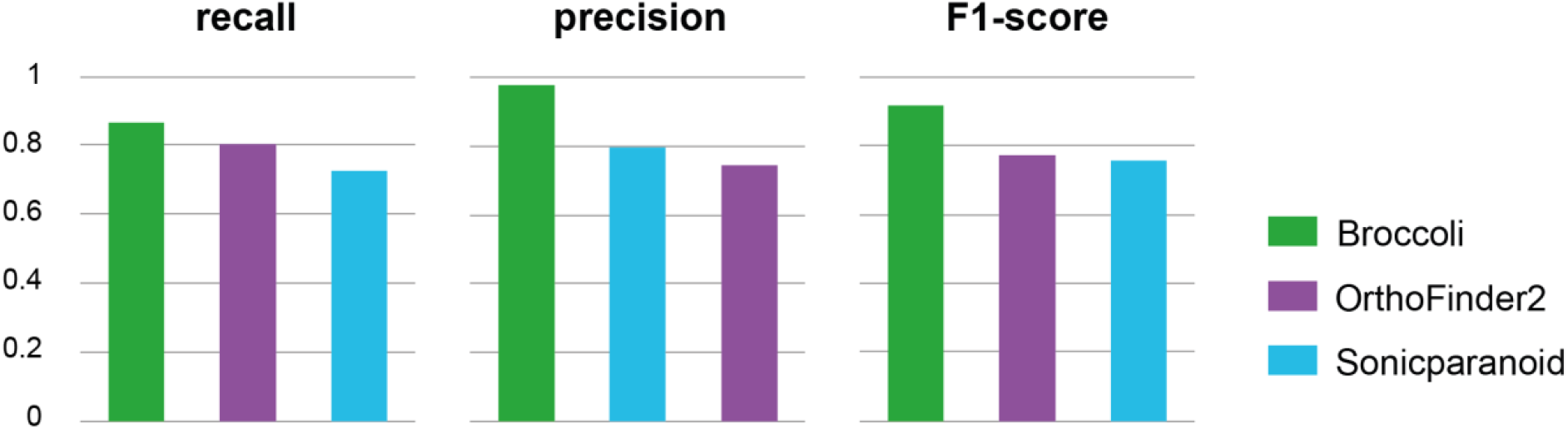
Benchmark of orthologous groups (paraBench dataset). Pipelines were ranked by their performance metric from left (highest value) to right (lowest value).

### Benchmark of orthologous pairs

We tested the orthologous pairs predicted by Broccoli, with its *r* threshold (Step 4) ranging from 0.3 to 0.7 in 0.1 increment (including the default value of 0.5), by using the large-scale Quest for Orthologs 2018 benchmark dataset (referred thereafter as QfO 2018 dataset) (Altenhoff, et al. 2016; Forslund, et al. 2017). This benchmark includes three groups of tests: (i) pairs of orthologs are compared to manually curated sets of gene phylogenies (using the TreeFam-A and SwissTree databases), (ii) the function of orthologs are compared to each other using the Gene ontology and Enzyme classification, assuming that orthologous genes should have the same function in different species, and (iii) the phylogenetic relationships of orthologous proteins are compared to reference species trees (the species Tree Discordance Benchmark, STDB, and its generalised version GSTDB), assuming that the gene trees should mirror the species trees. All benchmark values and scatter plots are available in Supplementary Material 3 and 4 respectively.

Orthologous pairs produced by Broccoli showed a strong agreement with the reference gene-phylogenies databases. When compared to the TreeFam-A database, only results produced by Broccoli were located on the Pareto frontier (Figure 3A), which is defines by the set of methods that are not outperformed by any other method in both recall (X-axis) and precision (Y-axis)(Altenhoff, et al. 2016). As expected, results corresponding to *r* = 0.7 (i.e. stringent threshold) scored the highest precision of all methods and results corresponding to *r* = 0.3 (i.e. relaxed threshold) were the most sensitive of all methods. In the case of the SwissTree database, a small reference dataset compared to the TreeFam-A database, all results were located on or closed to the Pareto frontier (Supplementary Material 4).

**Figure 3:**
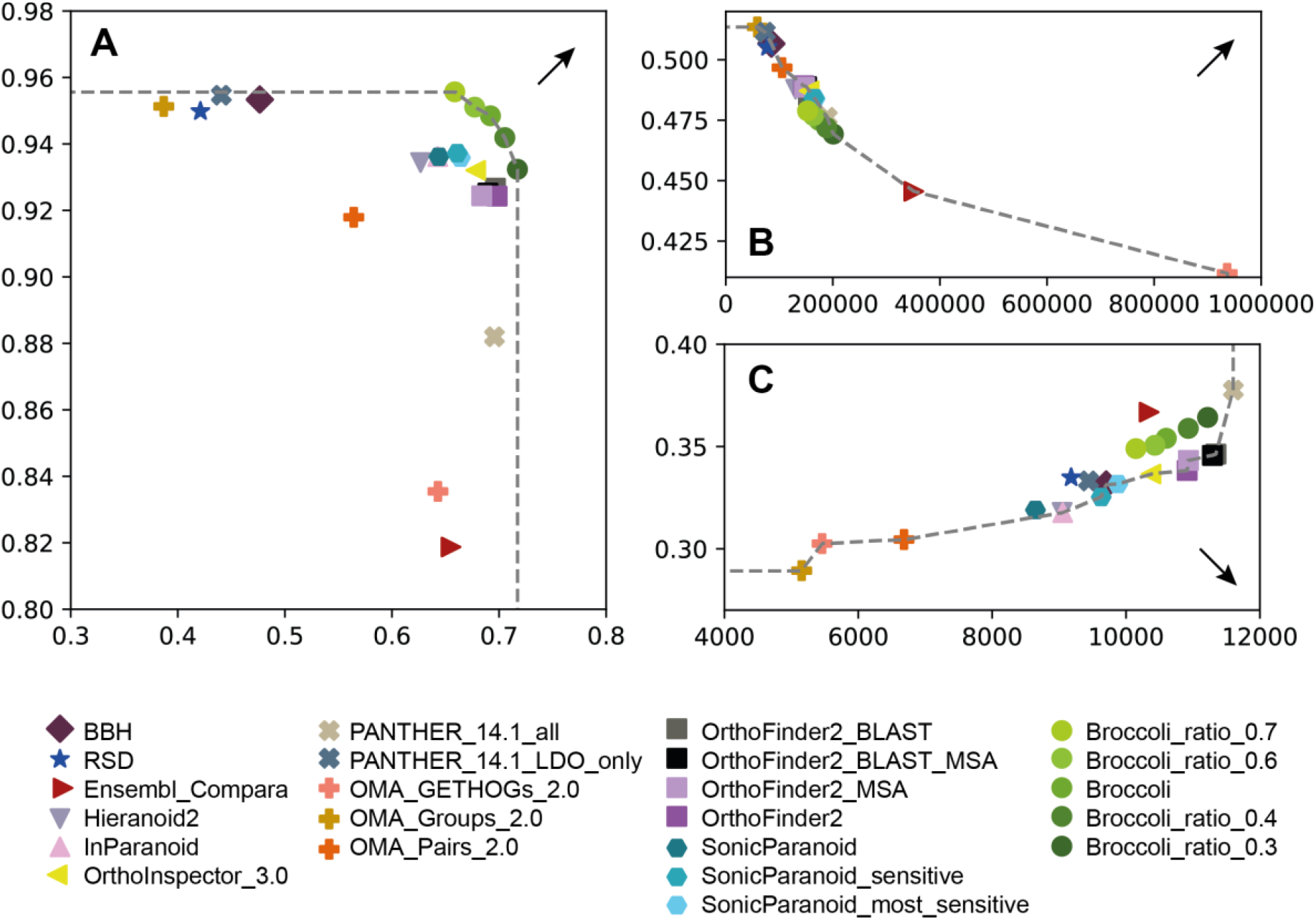
Benchmark of orthologous pairs (Quest for Orthologs 2018 dataset). The Pareto frontiers are represented by dotted lines and arrows indicate the direction toward better performances. In each of these three scatter plots, the X-axis and Y-axis correspond to a measure of recall and precision respectively. **A**: TreeFam-A benchmark (X-axis: true positive rate; Y-axis: positive predictive value). **B**: Geneontology benchmark (X-axis: number of orthologs; Y-axis: average Schlicker similarity). **C**: STD Fungi benchmark (X-axis: number of completed tree samplings; Y-axis: average Robinson-Foulds distance;).

In the two functional conservation tests, Broccoli results were similar to those of other methods (excluding two outlier methods, Ensembl Compara and OMA GEtHoG), with one (Gene ontology; Figure 3B) or two (Enzyme classification; Supplementary Material 4) Broccoli results on the Pareto frontier. As observed in the TreeFam-A benchmark, results corresponding to low *r* thresholds achieved higher precision (measured as the average Schlicker semantic similarity of functional annotations associated with orthologs) while results corresponding to high *r* thresholds showed higher sensitivity.

In contrast, the Broccoli performance in the species-trees discordance benchmarks are slightly worse than the ones of other methods: in most of these benchmarks, Broccoli results were distant from the Pareto frontier, with high sensitivity but high Robinson-Foulds distances (i.e. low precision) compared to other methods (e.g. STD Fungi in Figure 3C). The only exceptions were the STD Bacteria benchmark in which all Broccoli results were found on the Pareto frontier, and the GSTD Eukaryota benchmark in which two Broccoli results were found on the Pareto frontier (Supplementary Material 4).

### Chimeric proteins

In the absence of a specific gene-fusion benchmark, it is difficult to assess the quality of the predictions made by Broccoli. Nevertheless, a total of 1675 proteins were predicted to be the result of gene-fusion events from the QfO 2018 dataset (representing about 0.2% of all proteins). The number of chimeric proteins per species was highly heterogeneous, ranging from 0 (*Thermodesulfovibrio yellowstonii* and *Giardia intestinalis*) to 223 (*Zea mays*; Supplementary Material 3). Four species of this dataset showed particularly high numbers of chimeric proteins (namely *Z. mays*, *Phytophthora ramorum*, *Branchiostoma floridae* and *Monosiga brevicolis*). We hypothesize that these high prevalences are the consequences of errors in the gene prediction of these genomes. Broccoli was able to identify chimeric proteins that resulted from the combination of up to six orthologous groups and genes-fusions events shared by up to sixteen proteins. The latter case corresponds to the pentafunctional arom proteins present in multiple eukaryotic lineages (Richards, et al. 2006). However, Broccoli failed to recover some well-known gene fusion events such as the fusion of the dihydrofolate reductase and thymidylate synthase proteins present in ‘unikonts’ and absent in most ‘bikonts’ (Cavalier-Smith 2003), and the fusion of two tRNA synthetases shared by most metazoan species (Ray, et al. 2011).

### Running time analyses

Considering the large number of phylogenetic analyses performed by Broccoli (e.g. 658,421 phylogenies for the QfO 2018 dataset), it is expected to be several orders of magnitude slower than distance-based pipelines. We compared the runtimes of Broccoli to those of the two fastest distance-based algorithms Sonicparanoid and OrthoFinder2 (Emms and Kelly 2019) using datasets composed of 4 to 64 fungal proteomes. In these tests, the runtimes of Broccoli were found between those of Sonicparanoid and OrthoFinder2 (Figure 4). Regarding the two extremes of the speed spectrum, OrthoFinder2 with the MSA option was by far the slowest pipeline, and Sonicparanoid, which only performs half of similarity searches (Cosentino and Iwasaki 2019), was found to be the fastest for every dataset. The same speed rank was observed when analysing the QfO 2018 dataset, which contains 78 species, using 8 CPUs: Sonicparanoid (522 minutes), Broccoli (634 minutes) and OrthoFinder2 (850 minutes; we did not test the MSA option using this dataset).

**Figure 4:**
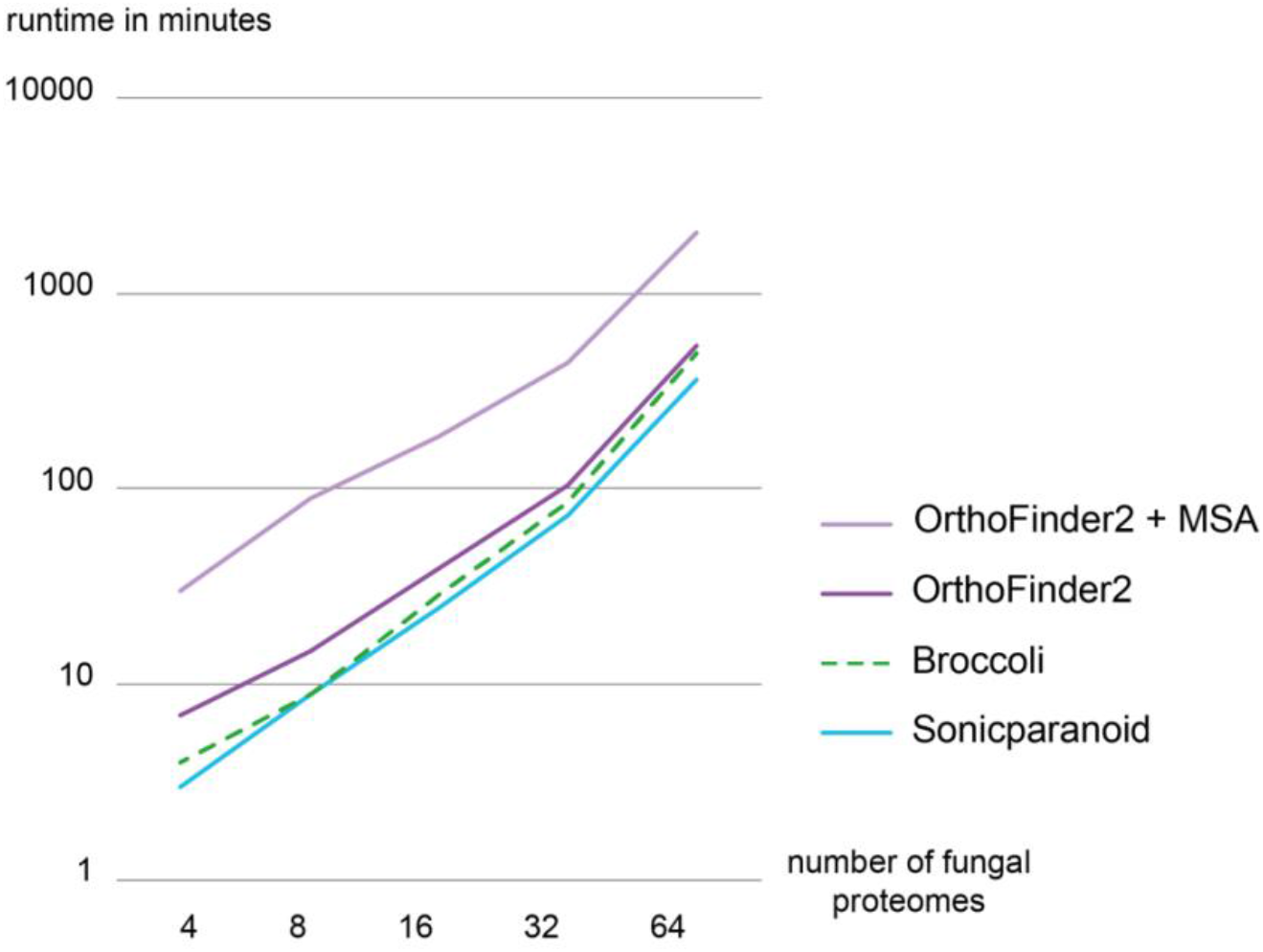
Efficiency tests. Runtimes, rounded to the nearest minute, were obtained using four CPUs and a dataset composed of 4 to 64 fungal species. Pipelines were ran using default parameters unless specified otherwise.

## Discussion

In this study, we introduced and tested a new phylogeny-based pipeline for orthology assignment. Since high-throughput phylogenetic analyses are challenging and time-consuming, the main idea behind Broccoli’s design is to perform ultra-fast phylogenetic analyses (i.e. pairwise alignments, simple trimming, NJ trees, midpoint rooting), and to rely on a performant community detection algorithm for the identification of relevant orthologous relationships. Broccoli has achieved this objective as it was found highly precise and sensitive on all tested benchmarks, with the noticeable exception of most QfO species tree benchmarks. While these specific benchmarks could possibly point to some limitations of the inferences made by Broccoli, it should be noticed that disagreements between gene trees and species trees are extremely common (e.g. (Marcet-Houben and Gabaldon 2009)), with many well-known sources of these discrepancies (e.g. incomplete lineage sorting, lateral gene-transfers, gene prediction errors; see introduction). Indeed, the high frequency of these discrepancies is the main reason as to why Broccoli employs a species overlap and not a species tree reconciliation approach for orthology delineation. Therefore, we believe that the use of distances between gene and species trees as a surrogate of precision measurement is questionable.

Finally, Broccoli showed high efficiency, with runtimes similar to those of the fastest distance-based pipelines, thanks to the parallelization of most tasks, an initial kmer clustering to simply proteomes, ultra-fast phylogenetic analyses and an efficient network analysis.

With a small subset of proteins being assigned to several orthologous groups, the clustering generated by Broccoli lies between classical gene classifications and protein domain subdivisions (e.g. Pfam database (El-Gebali, et al. 2019)). This fast and precise identification of chimeric proteins alongside their corresponding orthologous groups represents a promising avenue that should facilitate evolutionary studies of gene-fusion events (see also (Pathmanathan, et al. 2018)). However, future work is still required to widen the search of chimeric protein in the orthology network as the set of chimeric proteins currently identified by Broccoli appears to be incomplete.

Given the large variety of analyses performed by this pipeline (kmer clustering, phylogenetic analyses and network analysis), there are combinations of parameters that have not been tested, and parts that have not been fully optimised (e.g. trimming of the alignments, species overlap criteria). We are continuing to improve Broccoli by investigating parameters that should provide greater performances. In addition, its relatively high efficiency leaves much room for the implementation of more complex analyses. In its current form, Broccoli categorizes proteins (i.e. ortholog, chimeric) but does not infer evolutionary events (i.e. gene duplications, gene-fusions), which would require a reference species tree. We plan to implement an automatic species tree reconstruction using the supermatrix method (de Queiroz and Gatesy 2007), that will enable Broccoli to predict these evolutionary events as well.

## Acknowledgments

The authors thank Franz Lang and Luisa Orsini for their valuable comments on an earlier version of the manuscript. The authors also wish to thank Benjamin Buchfink for implementing new output fields in Diamond, and Toni Gabaldón for his seminal work on the phylogeny-based approach. Broccoli was developed as part of the DeepEuk collaborative project. Romain Derelle was supported by a NERC Highlight Topic grant (NE/N006216/1). John Colbourne was supported by UK NERC award Cracking the Code of Adaptive Evolution (NE/N016777/1).

## Notes

https://github.com/rderelle/Broccoli/

