## Supplementary Material 1 for "Broccoli: combining phylogenetic and network analyses for orthology assignment"

### Label propagation algorithm (LPA)

The LPA is applied independently to each connected component having a size superior to 3 nodes since it is not possible using this approach to infer several communities from a connected component of 3 or less nodes.

For each connected component, the implemented LPA works as follows:

1. the labels are initialized at all nodes (i.e. each node has a unique label).

2. nodes are ranked by their local clustering coefficient (decreasing order) and set to X.

3. for each node $x \in X$ chosen in that specific order, the node takes the label corresponding to the maximum value of S, calculated as:

$$S(l) = \sum_{i} w(xi)$$

where *i* represents the number of nodes connected to *x* and having the label *l*, and w(*xi*) represents the weight of the edge connecting *x* to *i* as described in the main manuscript. If several labels have an identical S value, tie is broken by choosing among these labels the one corresponding to the node having the highest edge weight w(*xi*).
If the new label *l* is different from the previous label of *x*, then it is considered as a change of label.

4. if at the end of the generation (i.e. after all nodes *x* of X had exchanged labels), there had been no observed change of label, the LPA stops. Otherwise, it goes back to (3) for a new generation.

This LPA is therefore:
_ asynchronous: labels are exchanged successively at step (3) instead of simultaneously at the end of the generation.
_ weighted: labels are chosen by summing the edge weights instead of simply summing the number of connected nodes.

Also, the original LPA involves random choices at step (2) to rank nodes, and at step (3) to break ties. Here, the LPA does not involve any random choice, which insures fast convergence of labels. Indeed, as shown in the figure below, labels reached convergence on the QfO 2011 dataset after only 2-3 generations in most connected components (data not shown).

### Detection of gene-fusions

The following figure illustrates the methodology implemented in Broccoli to identify gene-fusions among the network communities. Parameter names (i.e. arguments of step3 script) are coloured in red.

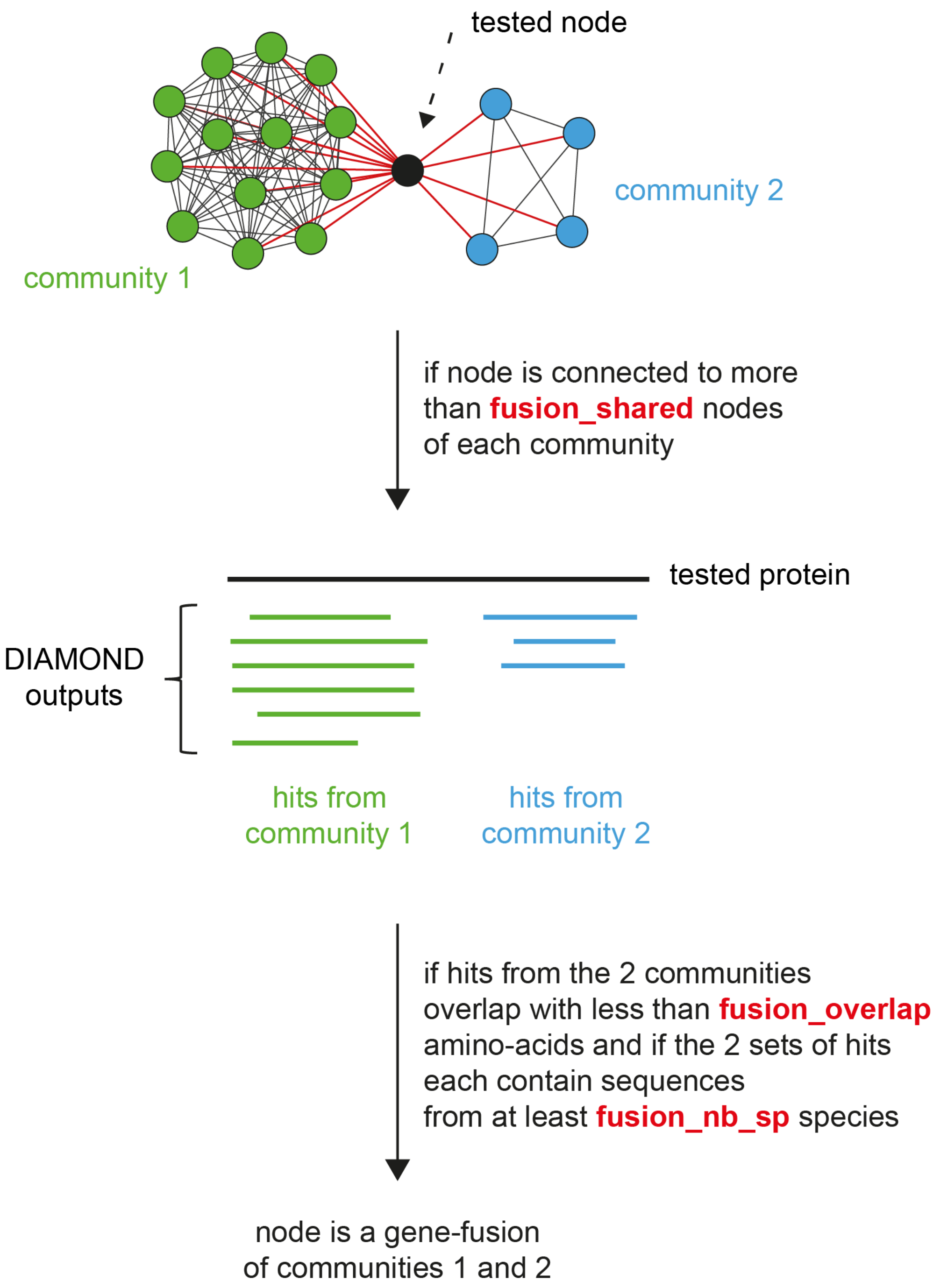
