## Supplementary Material 2 for "Broccoli: combining phylogenetic and network analyses for orthology assignment"

The only available benchmark dataset to assess the quality of orthologous group predictions, orthoBench [1], has a strong taxonomic bias toward vertebrates, with nine vertebrate species out of twelve bilaterian, and contains some errors (e.g. RefOG001 lacks all non-vertebrate proteins; personal observations). Therefore, we built an alternative benchmark dataset of comparable size to orthoBench with a taxonomic sampling more representative of standard analyses, named thereafter 'paraBench'.

The paraBench dataset was built from an in-house collection of phylogenomic markers [2, 3] (and unpublished work), most of which correspond to well characterised eukaryotic orthologous groups (OGs).
Phylogenomic markers were selected to be part of the paraBench dataset if they had at least one clear eukaryotic paralogous group (i.e. another eukaryotic OG branching as sister-group), so that the delineation of one OG could be easily made using the corresponding paralogous group. The selection of closely related OGs also allows to assess the precision of the different pipelines tested. Following to this criterion, a total of 52 OGs representing 14 gene families were selected.
17 proteomes from the QfO 2018 dataset were then chosen as representatives of most of eukaryotic supergroups and downloaded from EBI ftp server (ftp://ftp.ebi.ac.uk/pub/databases/reference_proteomes/QfO/ ; the list of species is provided below). These proteomes were BLASTed against the 53 selected alignments and hits were added to the alignments. We performed several rounds of phylogenetic analyses using Muscle version 3.8.31 [4] and RaxML version 8.2.12 [5] without alignment trimming to remove paralogs and only keep orthologs in the alignments, retaining as many divergent orthologs as possible.
The final reference classification contains 1357 proteins, and is similar in size to the orthoBench benchmark dataset:

| | orthoBench | paraBench |
|---|---|---|
| taxonomic level | animals | eukaryotes |
| number of species | 12 | 17 |
| number of OGs | 70 | 52 |
| number of proteins | 1695 | 1357 |

Precision and recall were calculated as follows:

$$\text{precision} = \frac{\text{TP}}{\text{TP} + \text{FP}} \qquad \text{recall} = \frac{\text{TP}}{\text{TP} + \text{FN}}$$

where TP, FP and FN correspond to true positives, false positives and false negatives respectively. Importantly, TP, FP and FN were only calculated among the 1357 proteins of the reference classification, insuring that ‘orthogroups’ produced by OrthoFinder [6] were not penalized by the addition of divergent in-paralogs not present in the reference classification.
Finally, F1-Score was calculated as the harmonic mean of precision and recall.

List of species included in the paraBench dataset:

*Homo sapiens*
*Nematostella vectensis*
*Monosiga brevicollis*
*Neurospora crassa*
*Ustilago maydis*
*Batrachochytrium dendrobatidis*
*Dictyostelium discoideum*
*Arabidopsis thaliana*
*Physcomitrella patens*
*Chlamydomonas reinhardtii*
*Paramecium tetraurelia*
*Plasmodium falciparum*
*Phytophthora ramorum*
*Thalassiosira pseudonana*
*Leishmania major*
*Giardia intestinalis*
*Trichomonas vaginalis*

<u>List of OGs included in the paraBench dataset:</u>

In the following pages, OGs were grouped by genes families, aligned with Muscle (default parameters) and phylogenetic trees were reconstructed using RaxML (command line: 'raxmlHPC-PTHREADS -f d -p 12345 -m PROTGAMMALG -T 8 -s ali_X.fa -n ali_X where 'ali_X.fa' is the name of the file containing the FASTA alignment of the gene family X) without alignment trimming. Each coloured box represents an OG.

#### Prohibitin

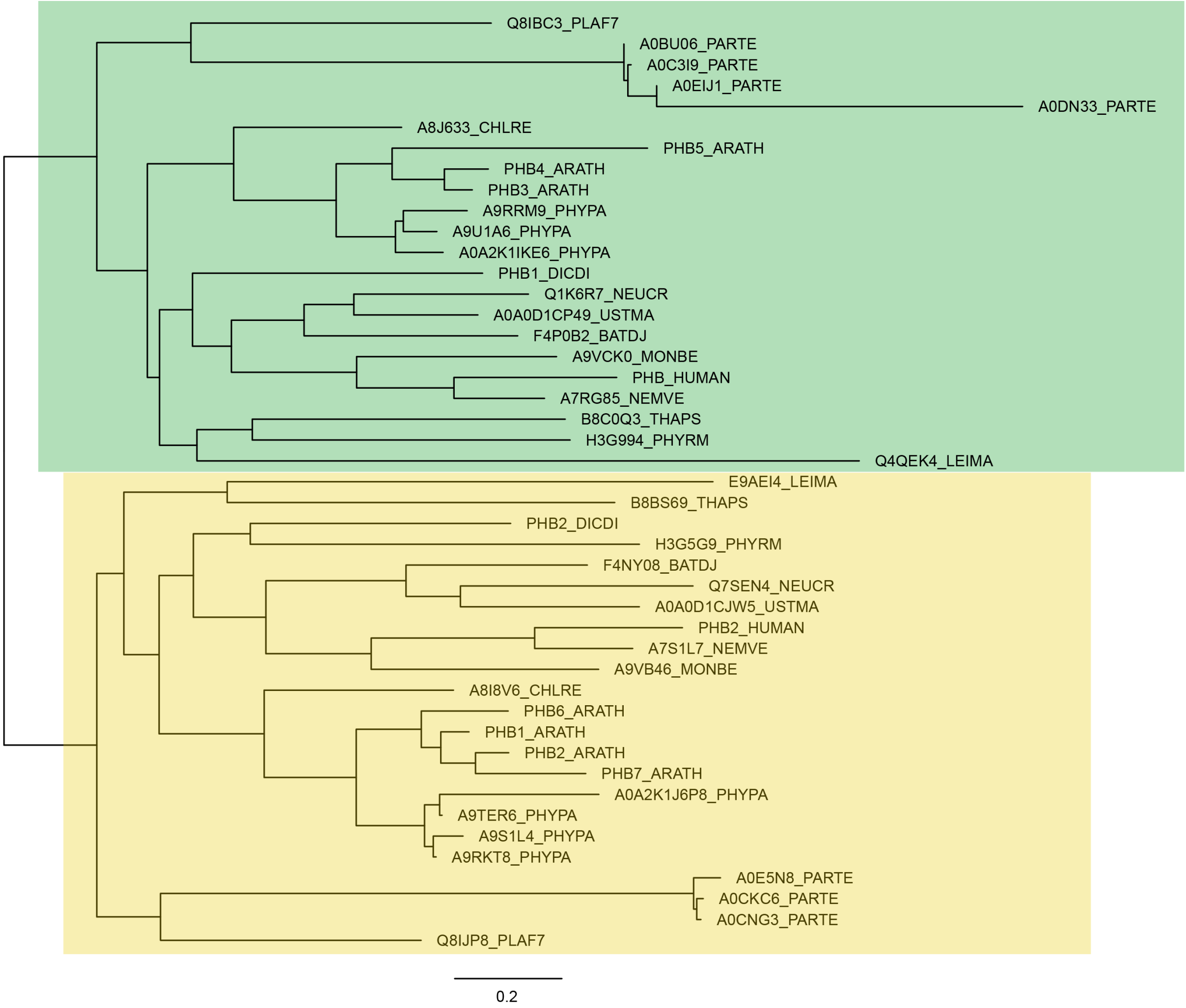

#### Nucleolar protein (nop)

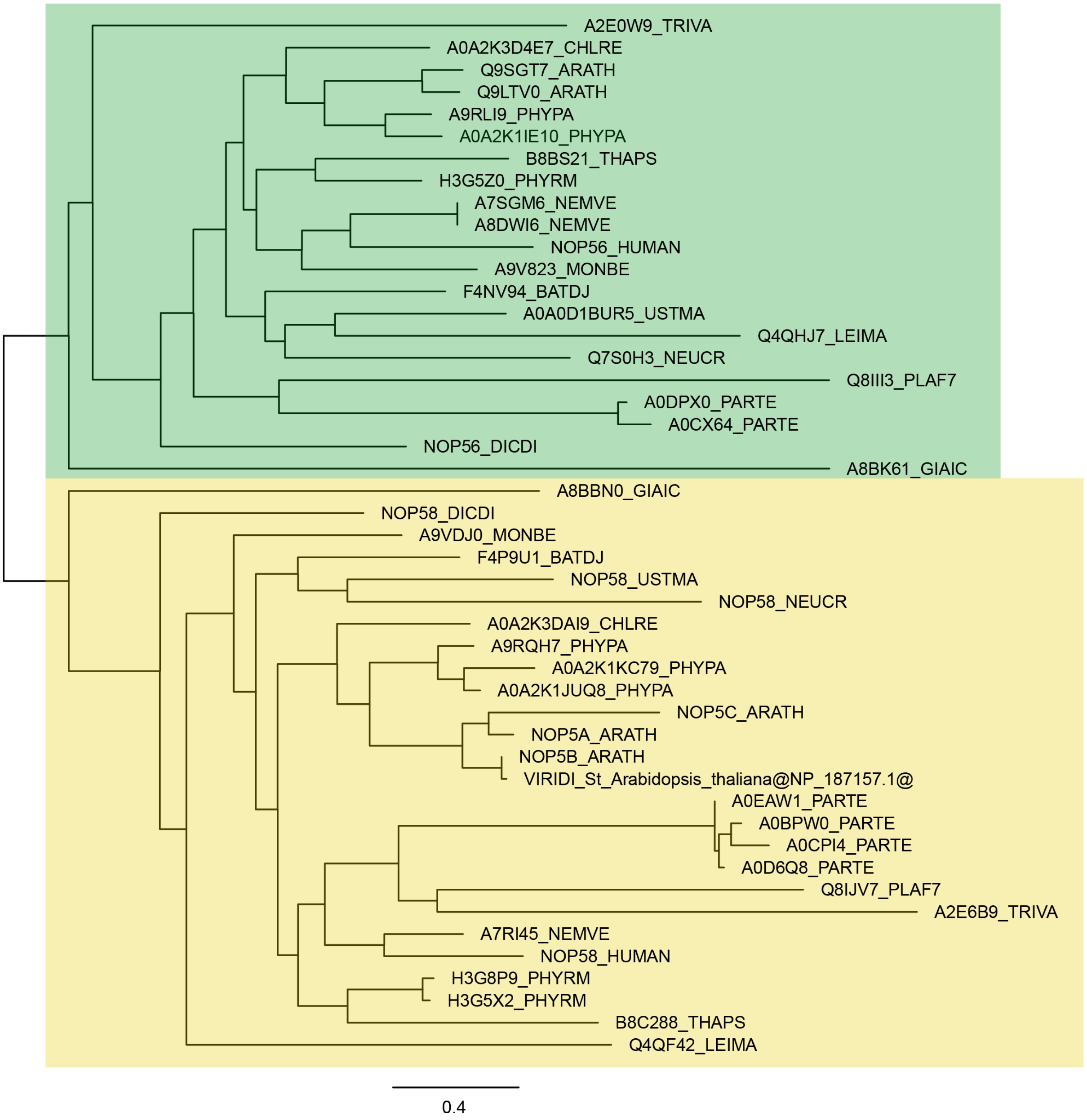

### Replication factor C (rfc)

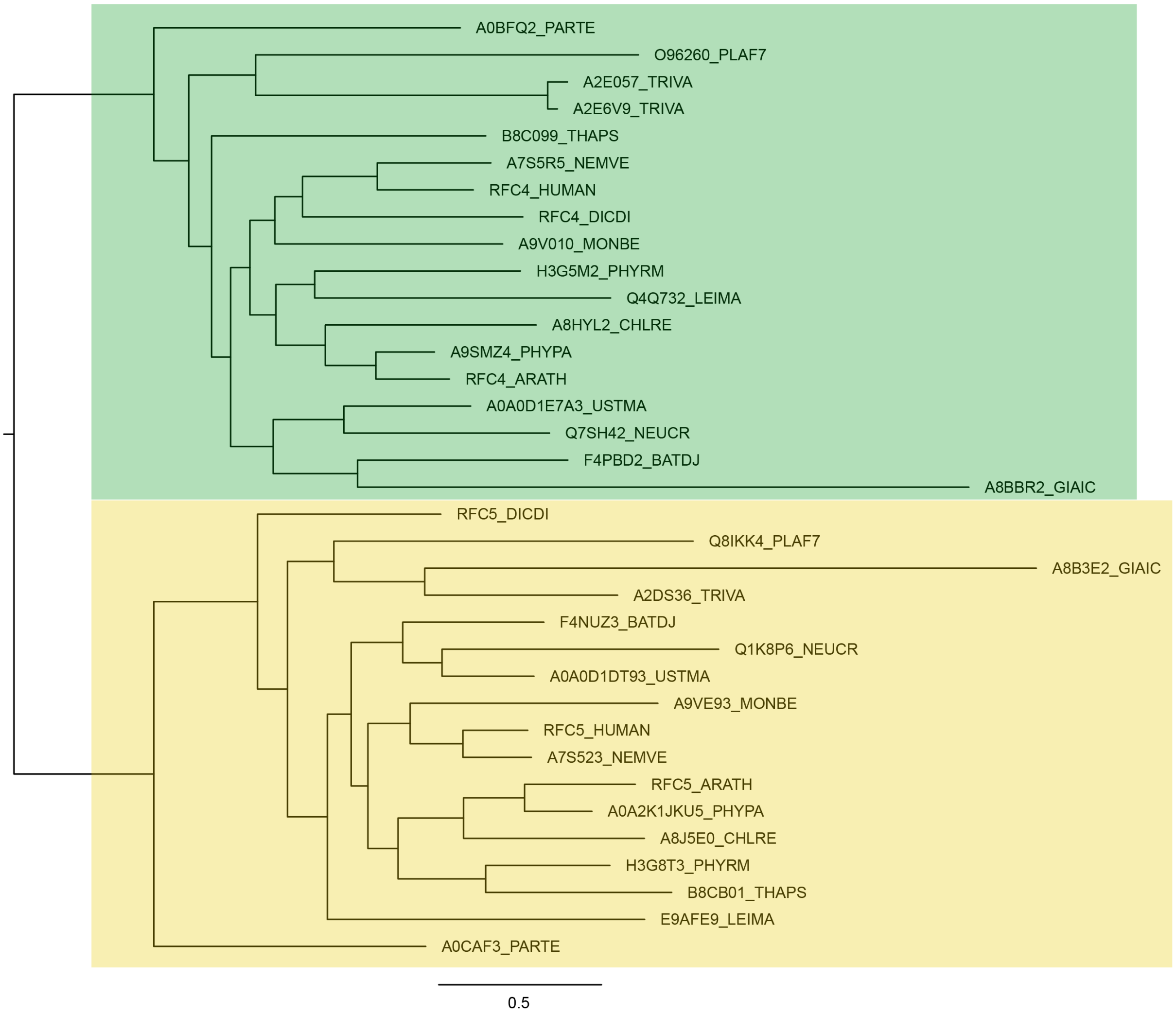

### Heat shock protein 70 (Hsp70)

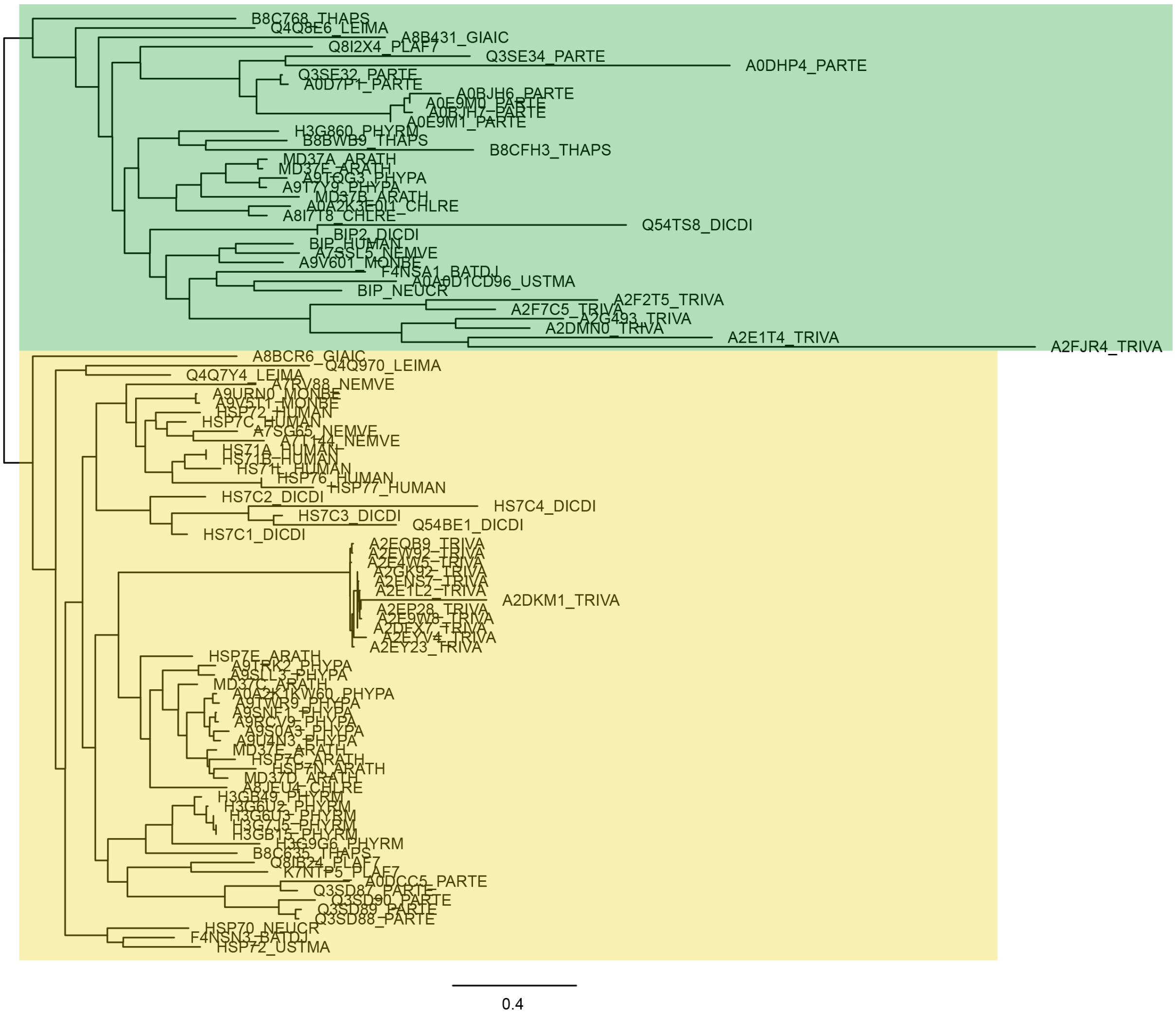

### Heat shock protein 90 (Hsp90)

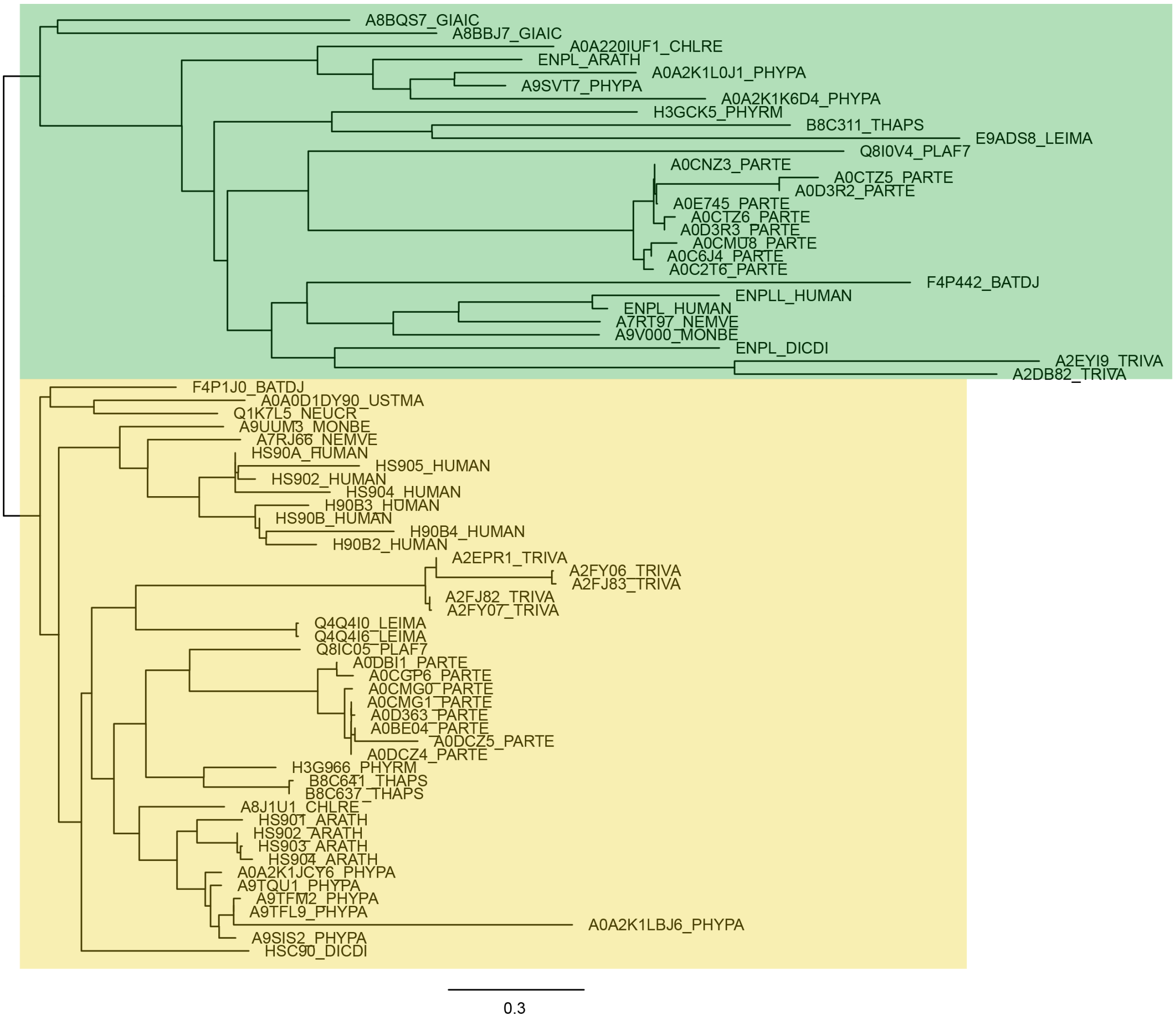

### developmentally-regulated GTP-binding protein (dGTP)

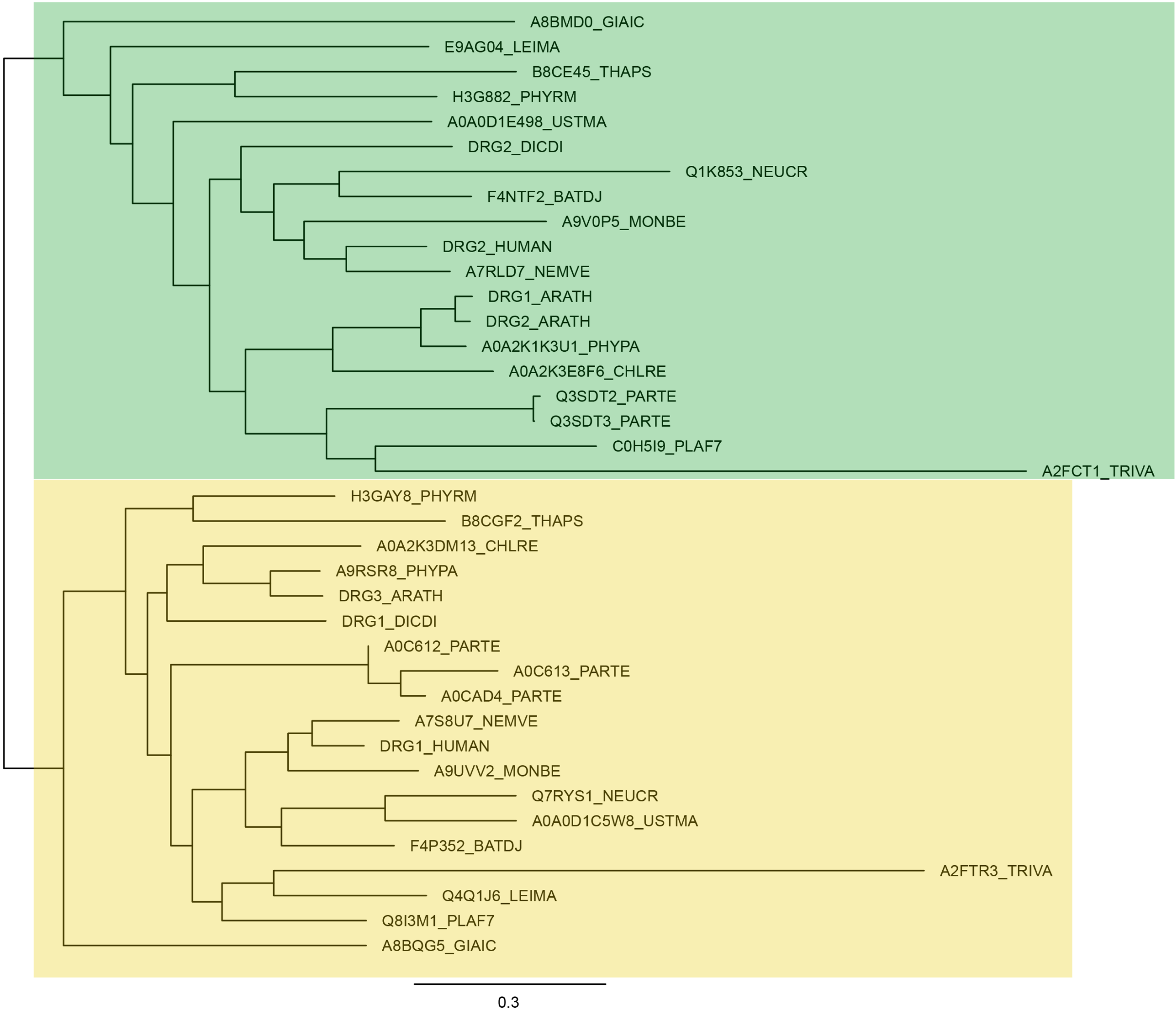

### Proteasome subunit alpha (psmA)

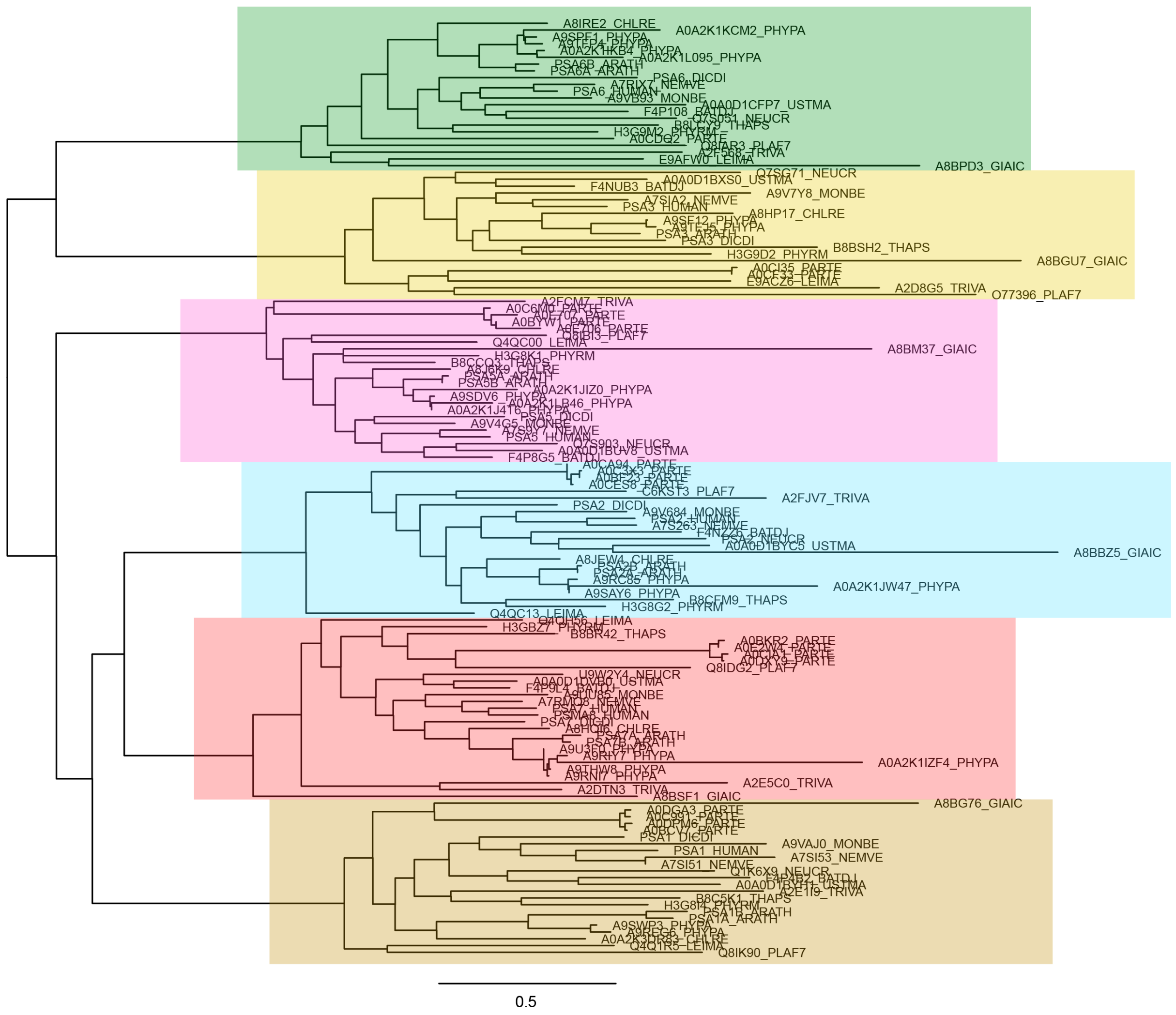

### Proteasome subunit beta (psmB)

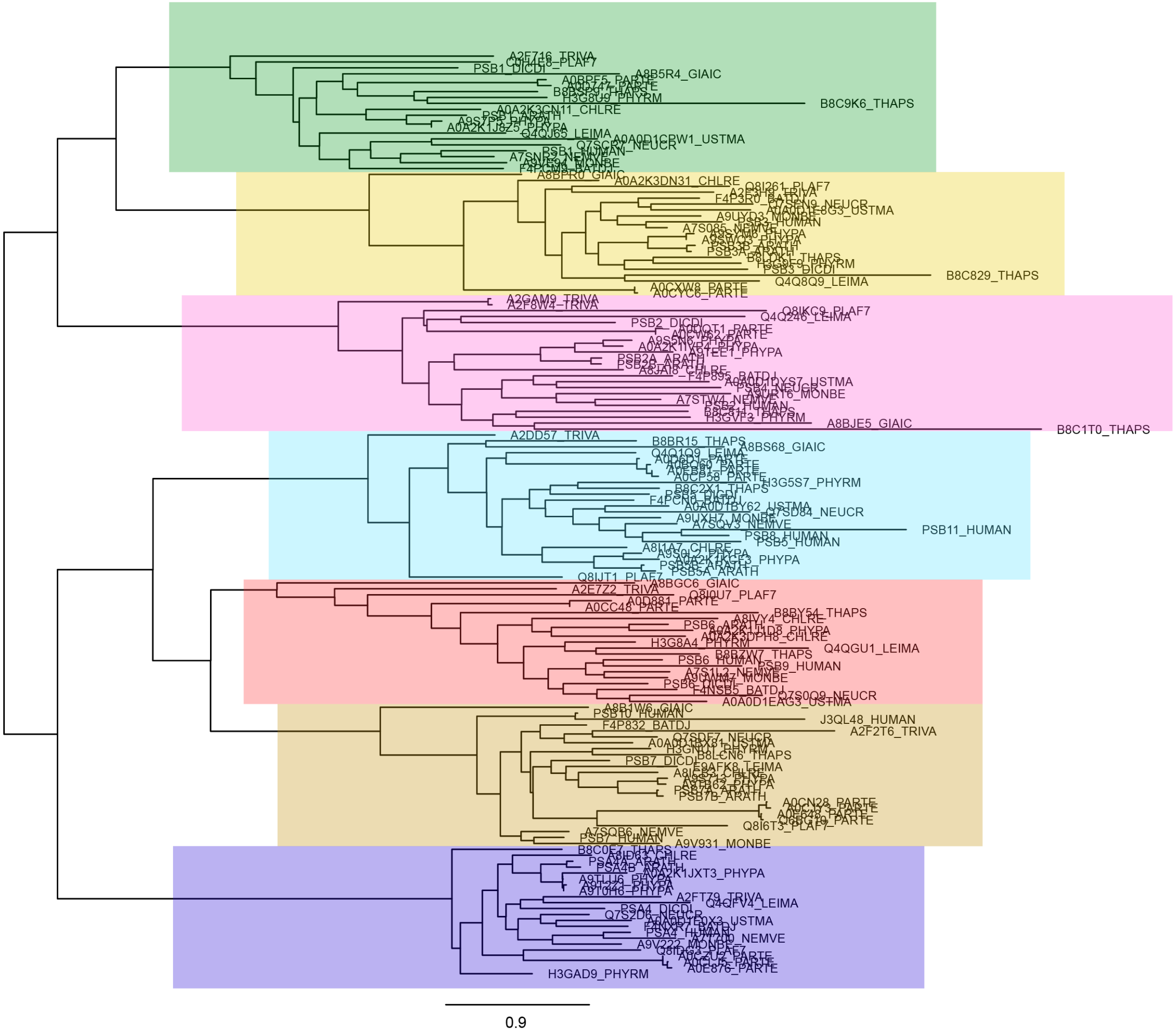

### AP complex subunit mu (apc)

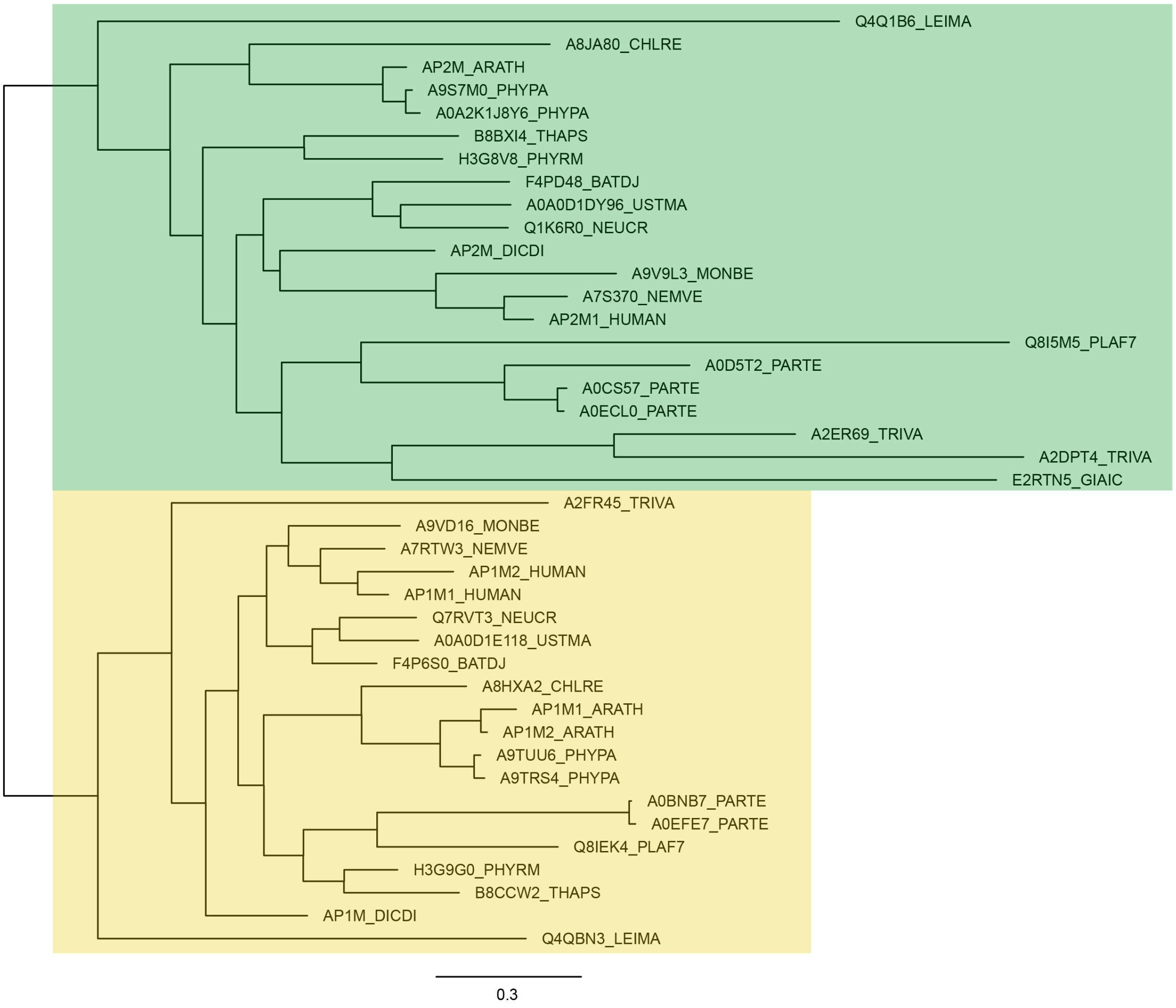

### Serine/threonine-protein phosphatase catalytic subunit (PP)

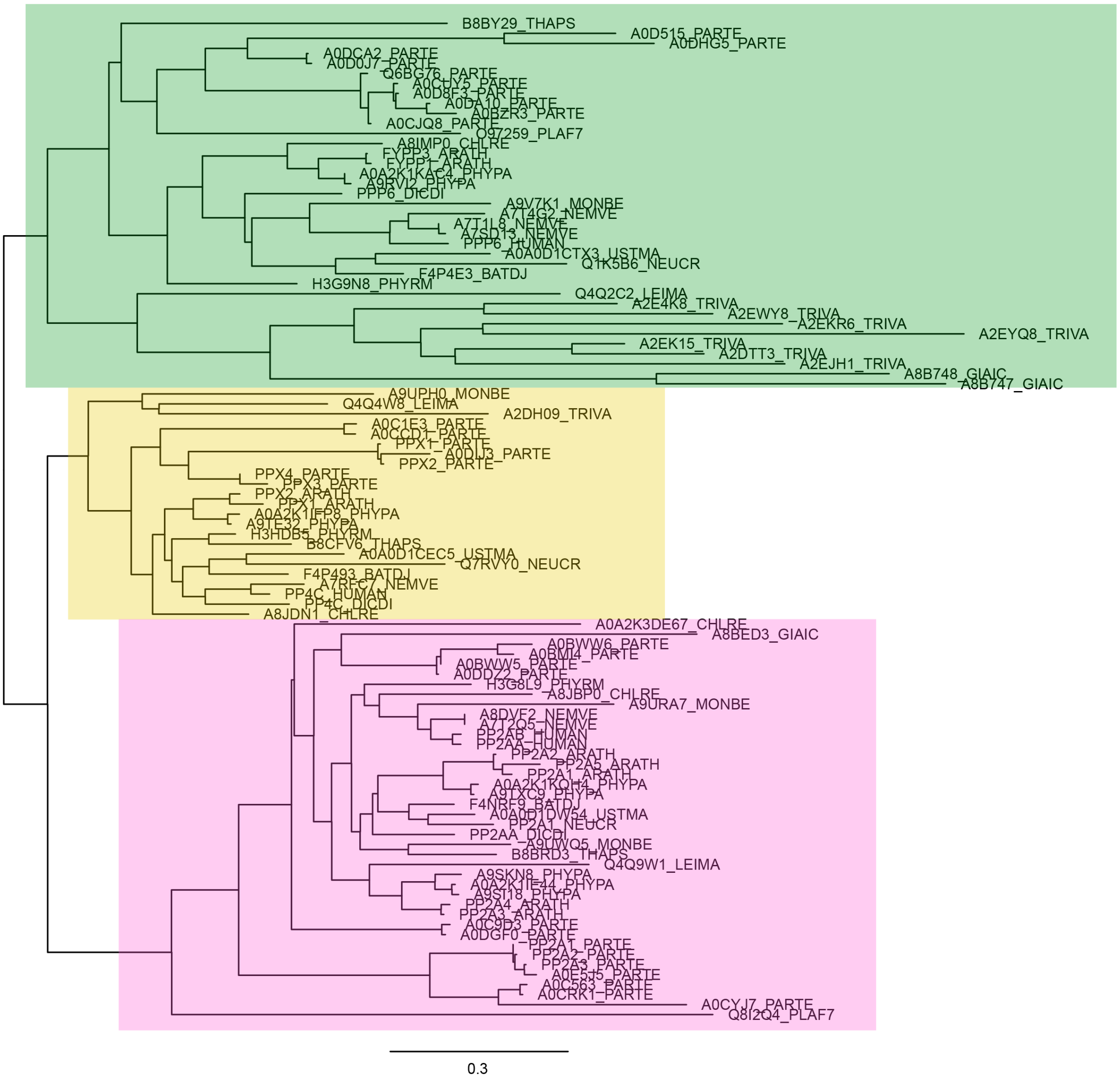

### ATP synthase subunit alpha (vat)

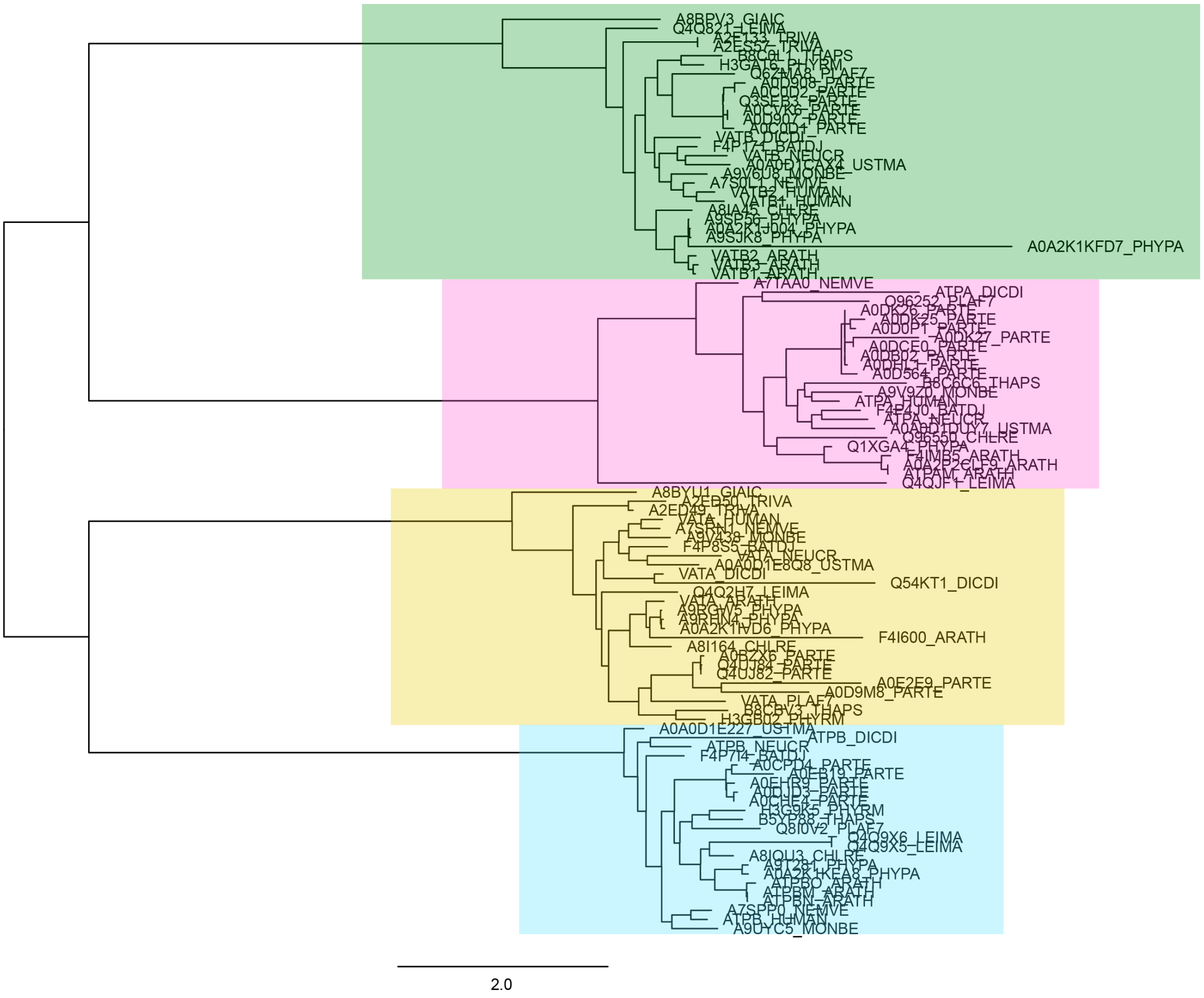

#### T-complex protein 1 (cct)

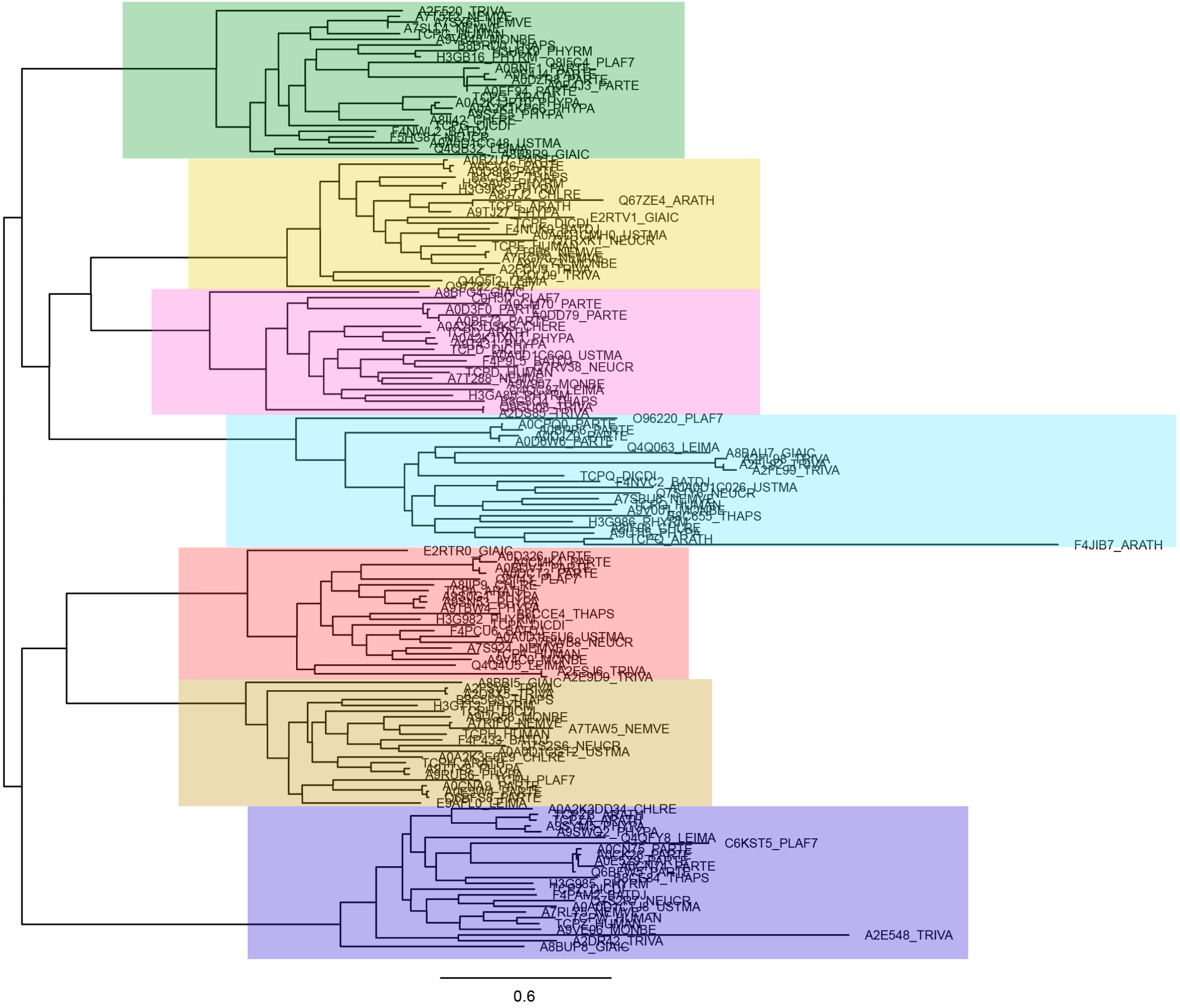

### DNA replication licensing factor (mcm)

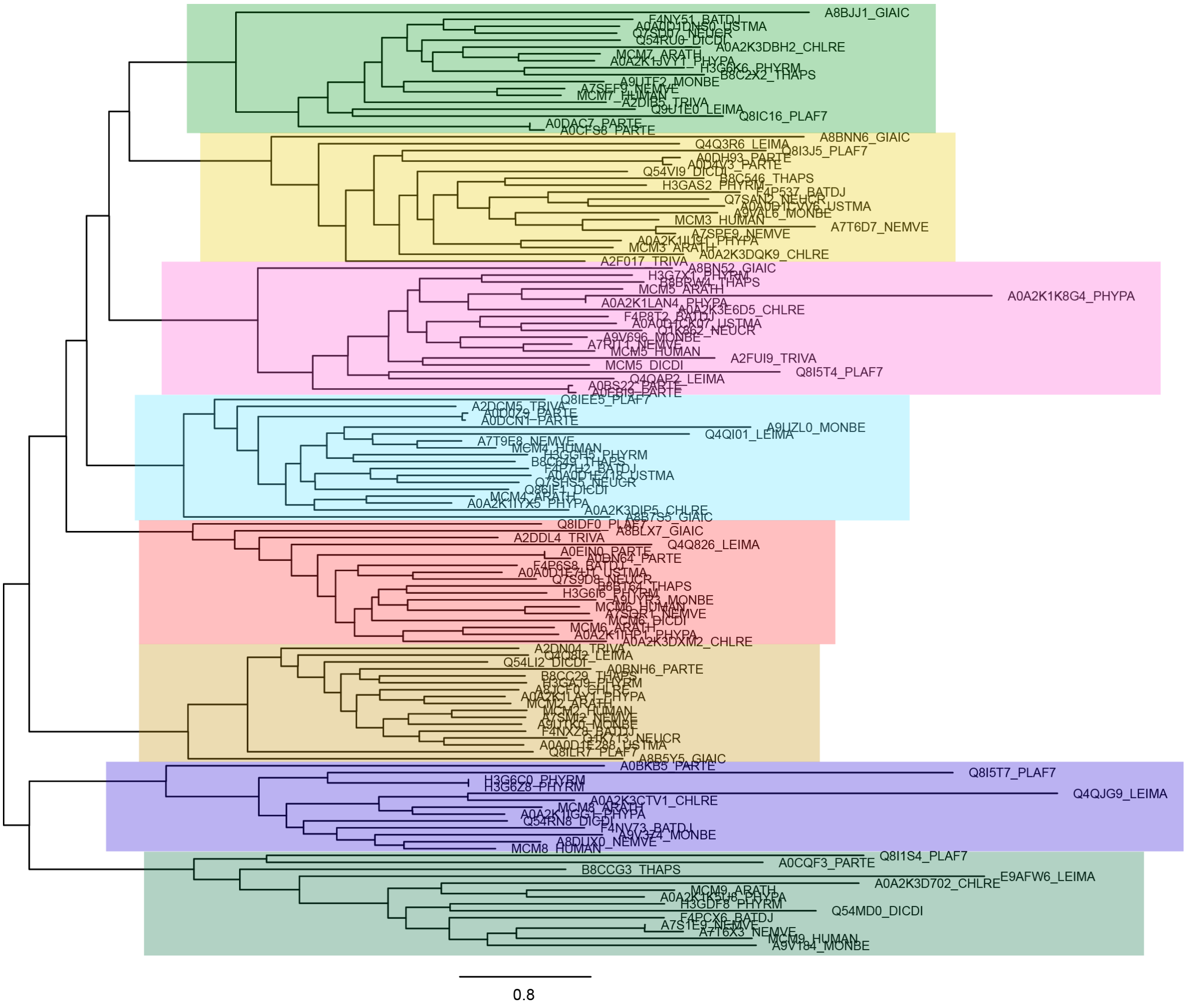

### Tubulin

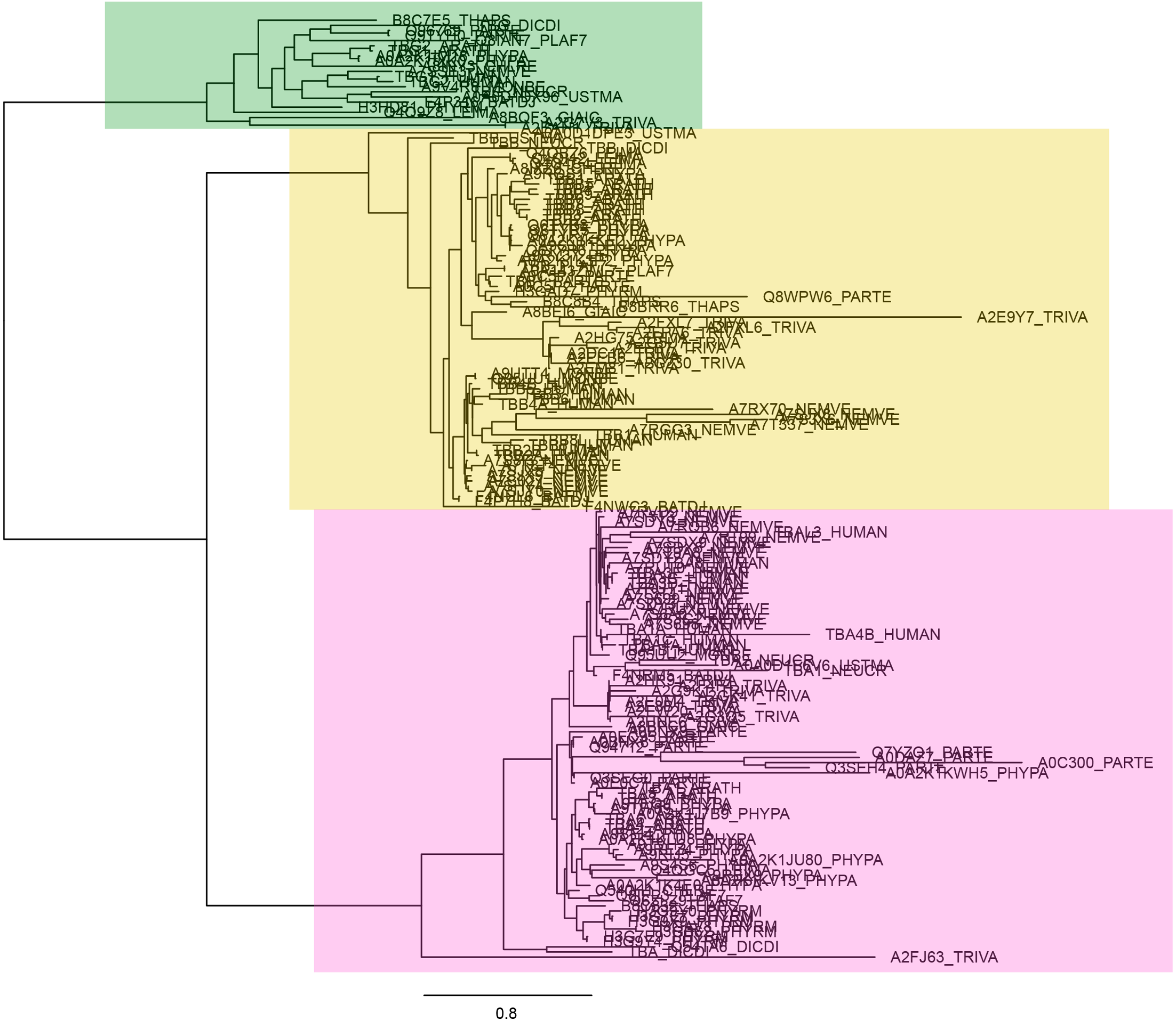

B8C7E5_THAPS
A0A2K1KWH5_PHYPA
Q8WPW6_PARTE
A2E9Y7_TRIVA
A7RX70_NEMVE
A7RCG3_NEMVE
A7T537_NEMVE
TBAL3_HUMAN
TBA4B_HUMAN
TBA1_NEUCR
Q7YZQ1_PARTE
A0C300_PARTE
Q3SEH4_PARTE
A0A2K1JU80_PHYPA
A2FJ63_TRIVA
0.8
