## Supplementary Material 4 for "Broccoli: combining phylogenetic and network analyses for orthology assignment"

The values corresponding to these plots are available in Supplementary Material 3, and the raw QfO 2018 results from which these values have been extracted are available in the compressed file deposited on.

### Gene trees benchmarks

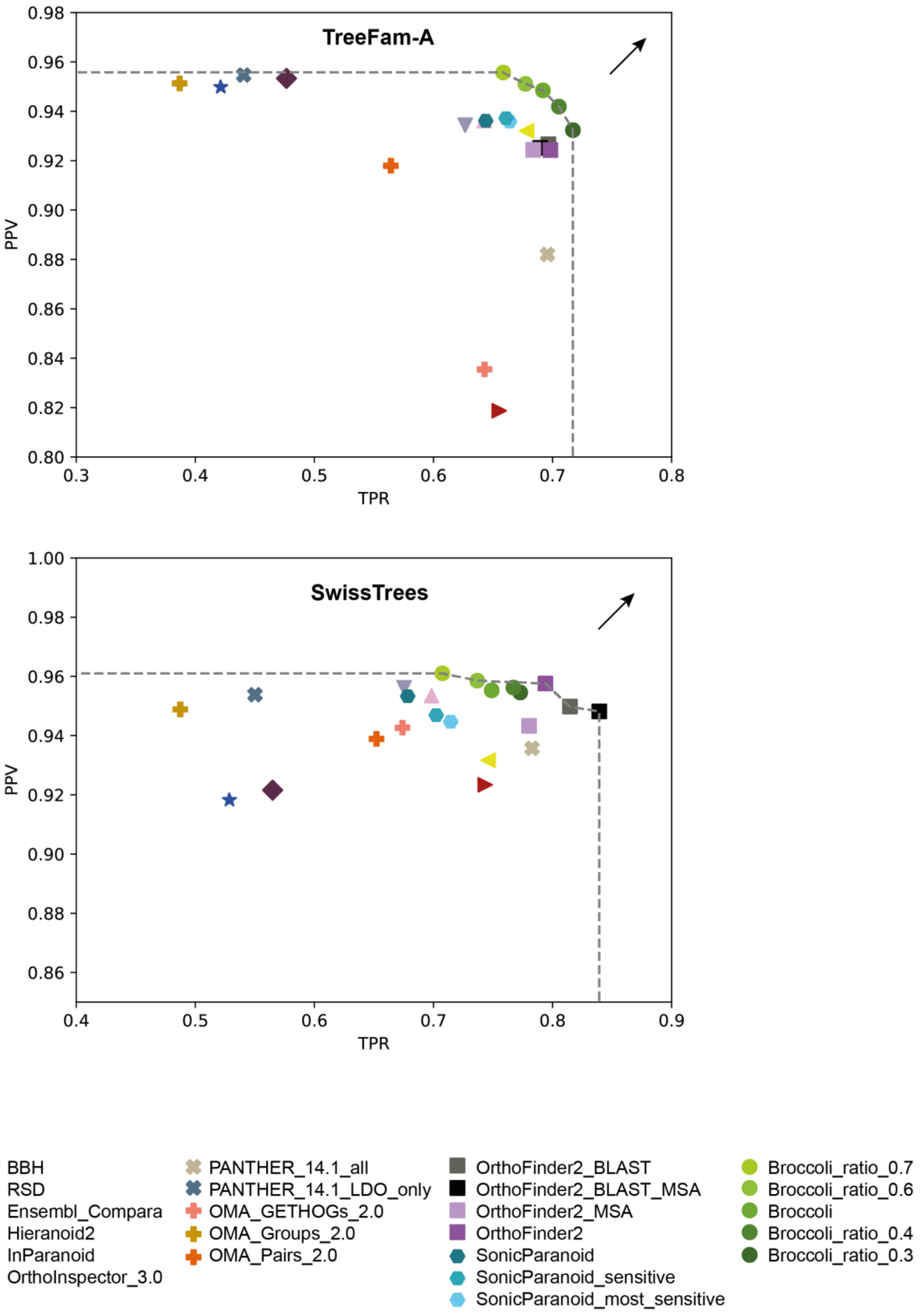

#### Function-based benchmarks

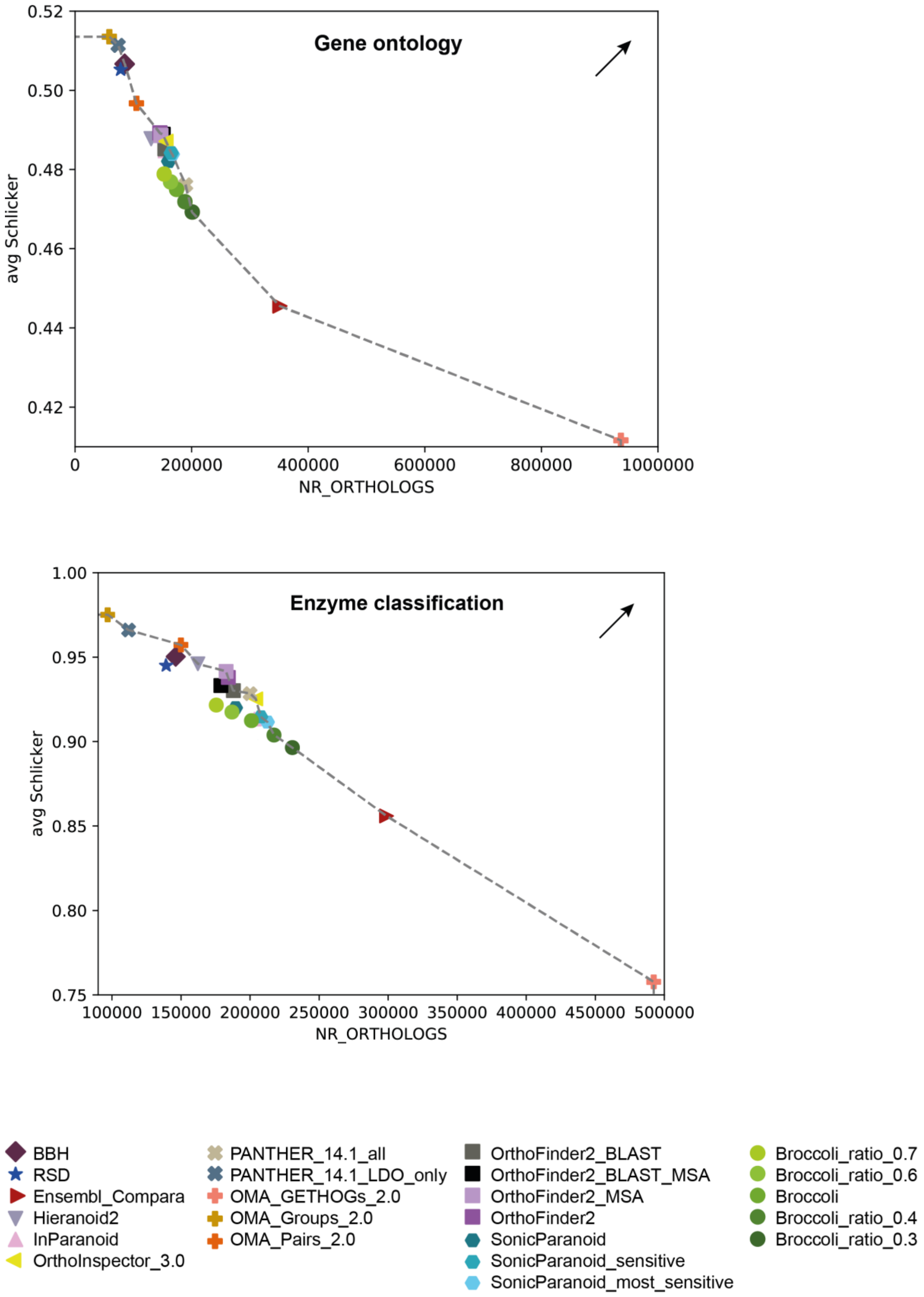

#### Species trees benchmarks

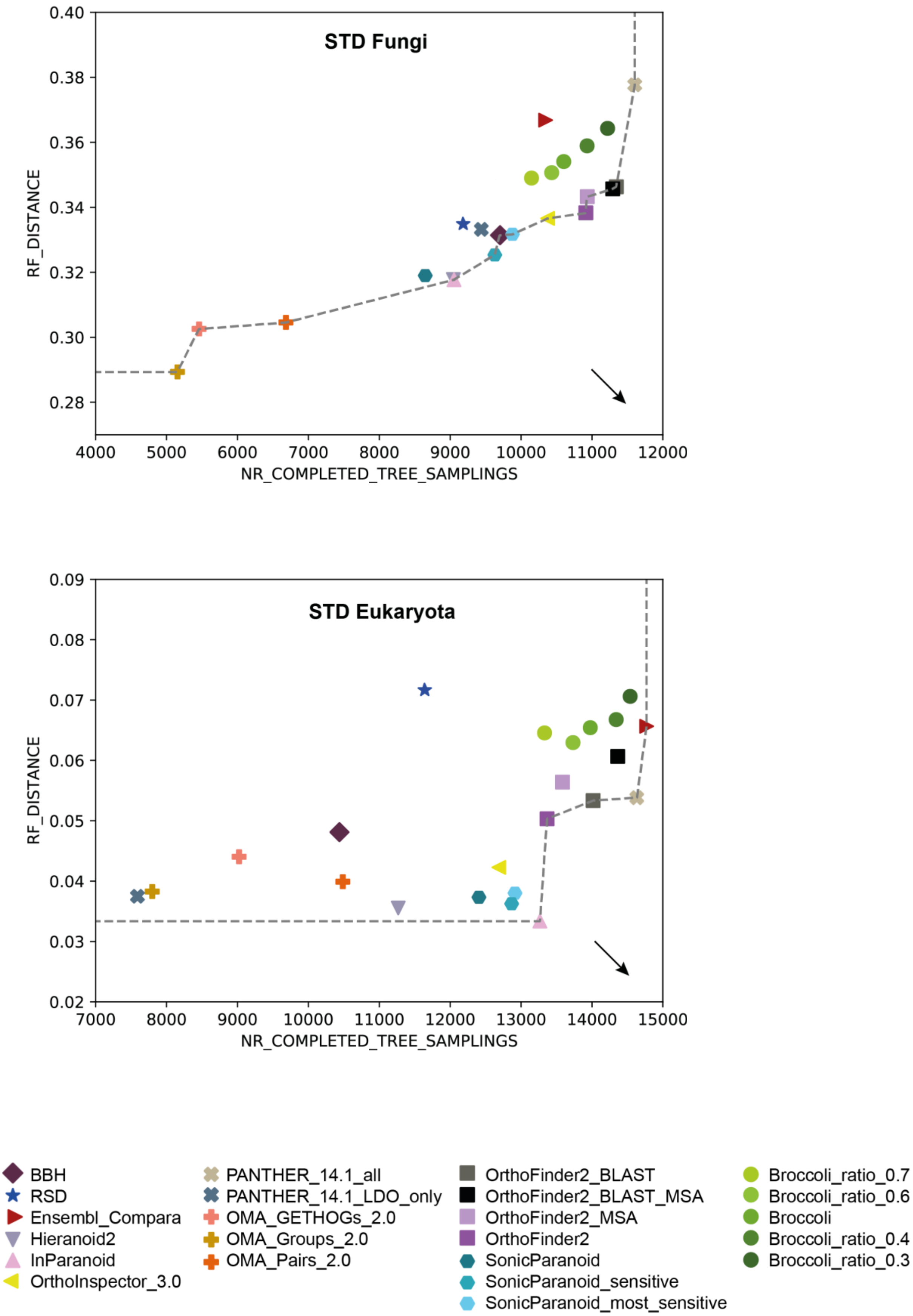

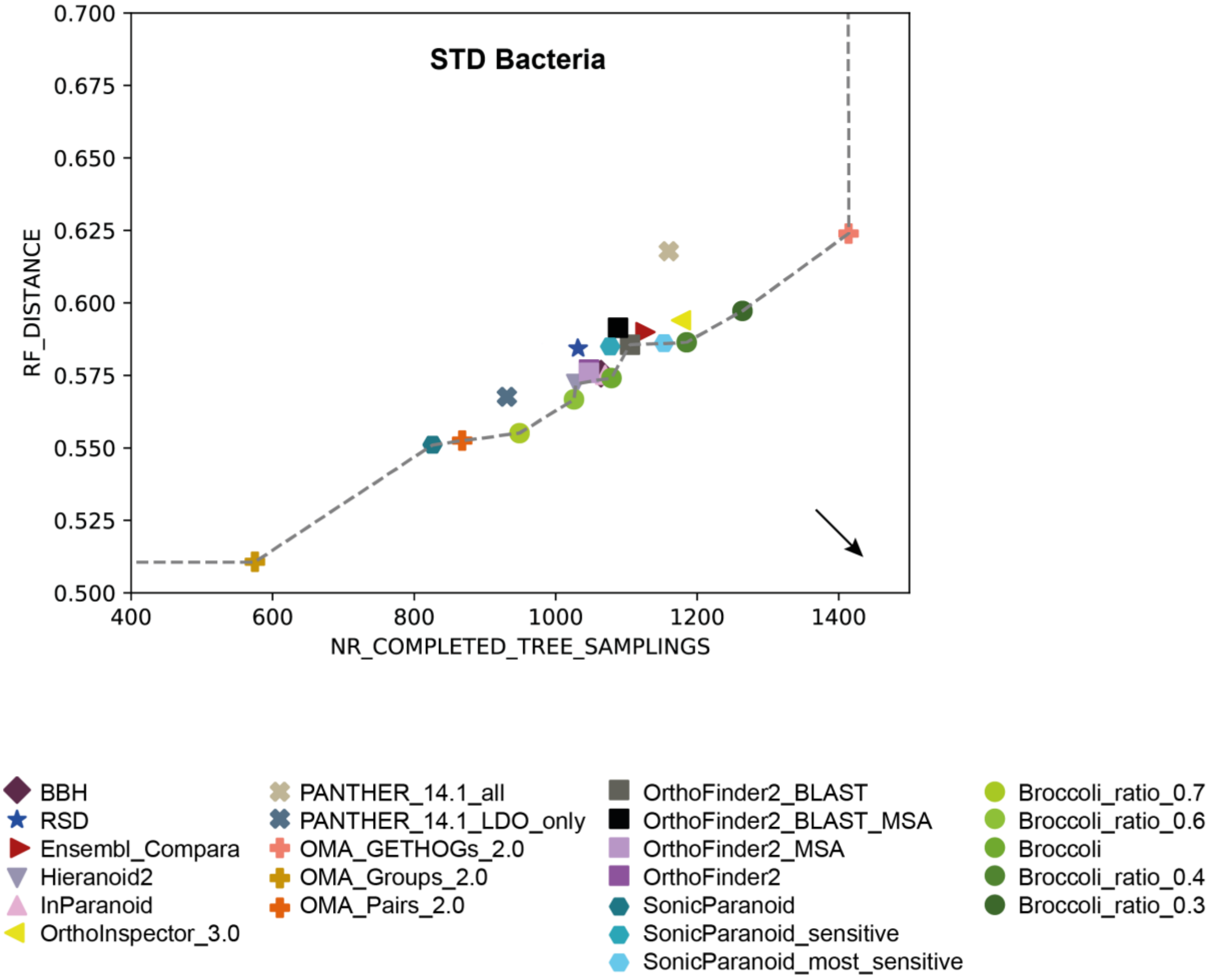
STD Bacteria
RF_DISTANCE
NR_COMPLETED_TREE_SAMPLINGS
0.700
0.675
0.650
0.625
0.600
0.575
0.550
0.525
0.500
400
600
800
1000
1200
1400
BBH
RSD
Ensembl_Compara
Hieranoid2
InParanoid
OrthoInspector_3.0
PANTHER_14.1_all
PANTHER_14.1_LDO_only
OMA_GETHOGs_2.0
OMA_Groups_2.0
OMA_Pairs_2.0
OrthoFinder2_BLAST
OrthoFinder2_BLAST_MSA
OrthoFinder2_MSA
OrthoFinder2
SonicParanoid
SonicParanoid_sensitive
SonicParanoid_most_sensitive
Broccoli_ratio_0.7
Broccoli_ratio_0.6
Broccoli
Broccoli_ratio_0.4
Broccoli_ratio_0.3

### Generalised species trees benchmarks

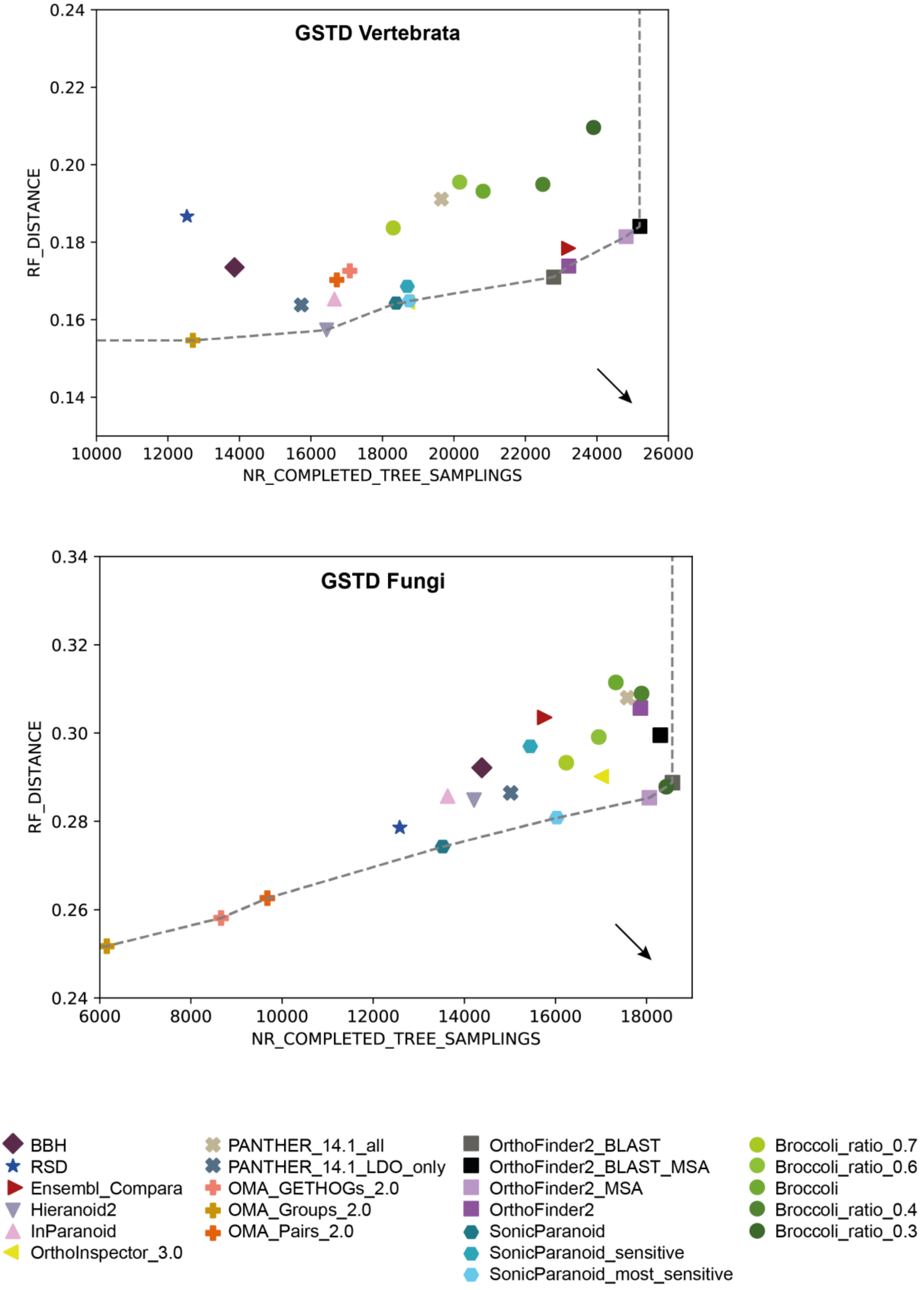

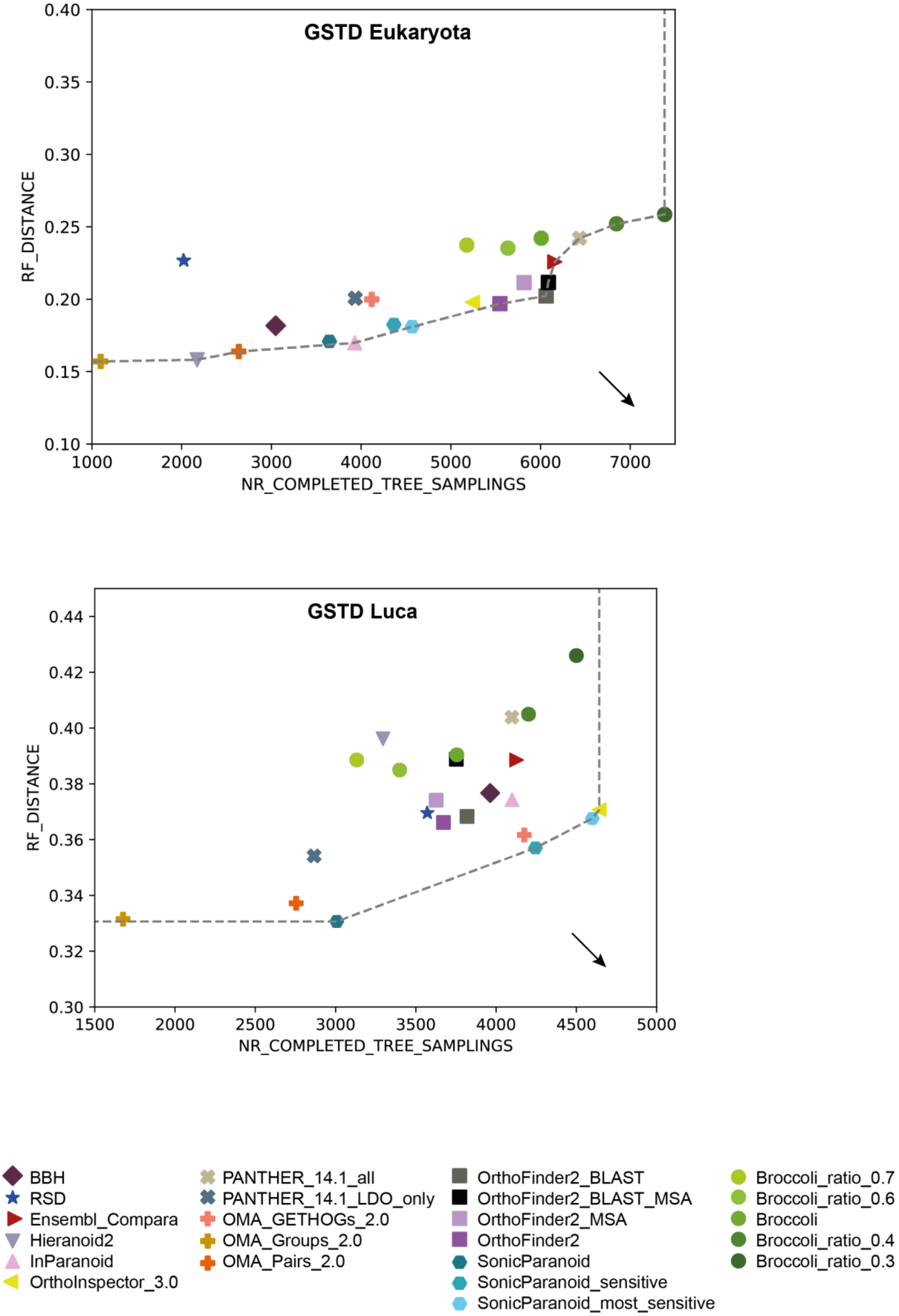
GSTD Eukaryota
RF_DISTANCE
NR_COMPLETED_TREE_SAMPLINGS
0.10
0.15
0.20
0.25
0.30
0.35
0.40
1000
2000
3000
4000
5000
6000
7000
GSTD Luca
RF_DISTANCE
NR_COMPLETED_TREE_SAMPLINGS
0.30
0.32
0.34
0.36
0.38
0.40
0.42
0.44
1500
2000
2500
3000
3500
4000
4500
5000
BBH
RSD
Ensembl_Compara
Hieranoid2
InParanoid
OrthoInspector_3.0
PANTHER_14.1_all
PANTHER_14.1_LDO_only
OMA_GETHOGs_2.0
OMA_Groups_2.0
OMA_Pairs_2.0
OrthoFinder2_BLAST
OrthoFinder2_BLAST_MSA
OrthoFinder2_MSA
OrthoFinder2
SonicParanoid
SonicParanoid_sensitive
SonicParanoid_most_sensitive
Broccoli_ratio_0.7
Broccoli_ratio_0.6
Broccoli
Broccoli_ratio_0.4
Broccoli_ratio_0.3
